# The free-living wellspring of symbiotic nitrogen fixation in *Bradyrhizobium*

**DOI:** 10.64898/2026.05.28.728359

**Authors:** Lu Ling, Sishuo Wang, Jinjin Tao, Marjorie Pervent, Kaitlyn E. Ho, Coline Sciallano, Alicia Camuel, Nico Nouwen, Eric Giraud, Haiwei Luo

**Affiliations:** Simon F. S. Li Marine Science Laboratory, School of Life Sciences and State Key Laboratory of Agrobiotechnology, The Chinese University of Hong Kong, Shatin, Hong Kong SAR; Department of Microbiology, Faculty of Medicine, The Chinese University of Hong Kong, Hong Kong SAR; PHIM Plant Health Institute of Montpellier, Université Montpellier, IRD, CIRAD, INRAE, Institut Agro, Montpellier, France; Department of Earth and Environmental Sciences, The Chinese University of Hong Kong, Shatin, Hong Kong SAR; Institute of Environment, Energy and Sustainability, The Chinese University of Hong Kong, Shatin, Hong Kong SAR

**Keywords:** *Bradyrhizobium*, symbiosis evolution, free-living nitrogen fixation, nitrogenase, genomic adaptation

## Abstract

The evolutionary origin of nitrogen-fixing symbiosis has been a long-standing question. To address this, we focused on *Bradyrhizobium*, a globally abundant bacterial genus that includes classic symbiotic lineages, which rely on the common Nod factor signaling pathway to form nodules, and close relatives capable of fixing nitrogen in a free-living state. We isolated 88 strains carrying the key genes for nitrogen fixation (*nif*) from non-legume environments and analyzed them alongside 586 public *Bradyrhizobium* genomes harboring these genes to reconstruct a robust phylogeny of *nif* genes. Analysis reveals that the earliest-diverging *nif* lineages are members capable of free-living nitrogen fixation, establishing this as the ancestral state. The Nod factor-dependent symbiotic lineages are polyphyletic, demonstrating at least three independent origins via horizontal acquisition of symbiosis islands. This evolutionary history is reflected in a genomic dichotomy: lineages capable of free-living nitrogen fixation possess a conserved *nif* island architecture that consistently includes the oxygen-protective gene *glbO*, whereas the symbiotic *nif*-associated regions are highly variable and universally lack *glbO*. Using both loss-of-function and gain-of-function genetic approaches, we show that *glbO* contributes significantly to nitrogenase activity under free-living conditions, whereas it is dispensable within the protected nodule environment. This work establishes a new framework for the evolution of symbiosis, identifying free-living ancestors as the source from which nitrogen-fixing symbiosis repeatedly and independently evolved in *Bradyrhizobium*.

**Significance:** For over a century, nitrogen-fixing bacteria called rhizobia have been celebrated for their symbiotic partnership with legumes like soybeans and peas. This study overturns the view that this symbiotic lifestyle was the original, advanced state. By discovering and analyzing diverse *Bradyrhizobium* bacteria from ordinary soils and non-legume plants, we show that the ancestor was capable of free-living nitrogen fixation. Root nodule nitrogen fixation evolved multiple independent times from free-living ancestors. We identify *glbO*, an oxygen-protective gene located immediately next to the nitrogen fixing genes, as critical for nitrogen fixation in variable soil conditions but dispensable and lost in *Bradyrhizobium* specialized inside nodules. This repositions free-living *Bradyrhizobium* as a major potential nitrogen source beyond legumes, with promise for sustainable agriculture.

## Introduction

Biological nitrogen fixation, the microbial conversion of inert atmospheric N_2_ into bioavailable ammonium, constitutes the primary natural nitrogen source for life on Earth (1–3). This process, carried out by diazotrophic prokaryotes or their derived organelle in eukaryotes, underpins global nitrogen cycling and ecosystem productivity (4–6). Among nitrogen-fixing microorganisms, root-nodule bacteria (rhizobia) are particularly notable for their capacity to establish intimate symbioses with leguminous plants through highly specific molecular exchanges (5, 7, 8). In the canonical interaction, rhizobia produce Nod factor (NF) signals that are perceived by the host plant and trigger root hair infection, culminating in the formation of a specialized symbiotic organ, the root nodule. This well-established paradigm is not universal: two NF-independent symbiotic pathways have also been identified, which differ in their dependence on a bacterial type III secretion system (T3SS) (9, 10). Collectively, these diverse symbioses provide plants with fixed nitrogen in exchange for carbon and form a cornerstone of sustainable agriculture (11).

For decades, the evolution of nitrogen fixation in rhizobia has been interpreted through a legume-centric lens (12–14). Studies of model genera such as *Rhizobium*, *Sinorhizobium,* and *Mesorhizobium* often portrayed the symbiotic lifestyle as an evolutionary pinnacle (12, 15, 16). In these models, symbiotic capability is frequently acquired via horizontal transfer of “symbiosis islands”, mobile genetic elements carrying essential *nod* (nodulation) and *nif* (nitrogen fixation) genes (13, 14, 17). Consequently, free-living nitrogen-fixing strains have been considered derived outliers.

This prevailing paradigm is challenged by the genus *Bradyrhizobium*, a globally abundant and ecologically versatile group of soil bacteria with a major influence on nitrogen budgets (18, 19). Although it contains classic symbiotic models like *B. diazoefficiens* USDA110 and *B. elkanii* USDA61 (10, 20), its ecological role extends beyond legume symbiosis. Accumulating evidence indicates that *Bradyrhizobium* lineages carrying *nif* genes and capable of free-living nitrogen fixation (defined here as “FL*nif*”: *nod*-free but *nif* carrying) are prevalent in soils worldwide (21–23), not mere outliers. Some FL*nif* strains belong to the *nod*-free but nodulating *Bradyrhizobium* supergroup (traditionally called Photosynthetic *Bradyrhizobium* or PB supergroup). This group lacks *nod* genes but forms nitrogen-fixing nodules with some tropical legumes of the genus *Aeschynomene* via an alternative, genetically undefined process likely encoded in its core genome (23). Furthermore, these FL*nif* members form endophytic associations with non-leguminous plants, such as rice, maize, sugarcane, and sweet potato, potentially supplying fixed nitrogen (23–30). In niches like the rhizosphere, they can constitute a significant portion of the diazotrophic community (31, 32), suggesting a substantial contribution to environmental nitrogen fixation.

The evolutionary history of *nif* genes within *Bradyrhizobium* has remained unresolved, primarily because insufficient genomic data precluded a robust *nif* gene phylogeny. This lack of resolution leaves fundamental questions unanswered. Four competing scenarios for the origin of NF-dependent symbiotic *nif* clusters remain indistinguishable: (1) vertical inheritance from a single common symbiotic ancestor of the entire genus, followed by loss in lineages capable of free-living nitrogen fixation; (2) independent horizontal acquisition of symbiosis islands by distinct free-living ancestors; (3) a single origin from a free-living ancestor, followed by horizontal transfer and radiation of the symbiotic *nif* cluster within *Bradyrhizobium*; or (4) horizontal acquisition of the symbiotic *nif* cluster from other rhizobial genera, such as *Rhizobium*, *Sinorhizobium*, or *Mesorhizobium*. This knowledge gap is directly constrained by the scarcity of sequenced free-living diazotrophic *Bradyrhizobium*, limiting phylogenetic reconstruction power (22, 33). Additionally, comparative genomics suggests adaptations such as the consistent presence of the oxygen-protection gene *glbO* (encoding a truncated hemoglobin for oxygen protection) in free-living *nif* clusters and absent in symbiotic ones (22), but direct functional evidence has been lacking.

This study addresses these questions by expanding the known genomic diversity of free-living nitrogen-fixing *Bradyrhizobium* and integrating them into an evolutionary and functional framework. We isolated and sequenced 88 FL*nif* strains from non-legume ecosystems, providing unprecedented resolution. Using this expanded dataset, we reconstruct the evolutionary history of *nif* genes to test hypotheses on the origin of symbiosis. We performed comparative genomic analysis to identify features of *nif* islands correlated with lifestyle. Finally, we used genetic manipulation and phenotypic assays to test the functional role of *glbO* in the *nif* island and establish a hierarchy of nitrogen fixation efficiency across *Bradyrhizobium* diversity.

## Results

### Genomic diversity of Bradyrhizobium capable of free-living nitrogen fixation

Available isolates have constrained our understanding of *Bradyrhizobium* that carry *nif* genes and fix nitrogen in a free-living state (FL*nif*). To address this gap, we isolated and sequenced 88 FL*nif* strains from roots and surrounding soils of six non-legume plants across three geographic regions in China: northeast (Liaoning), central (Hunan), and southeast (Hong Kong) (**Fig. S1, Dataset S1**). Host plants included rice (*Oryza sativa* subsp. *indica* and *japonica*), *Excoecaria agallocha*, *Camphora officinarum*, *Houttuynia cordata*, and *Phragmites australis*, representing cropland, forest, grassland, and wetland ecosystems (**Fig. S2, Dataset S1**). Isolation attempts from maize in Shanxi and Anhui provinces were unsuccessful (**Fig. S1, Dataset S1**).

Phylogenomic analysis confirmed these new isolates substantially broaden the known diversity of *Bradyrhizobium* capable of free living nitrogen fixation. They span five of seven recognized supergroups: the *nod*-free but nodulating (PB), Kakadu, *B. jicamae*, *B. japonicum*, and *B. elkanii* supergroups (**Fig. 1A**). Our efforts yielded FL*nif* representatives from supergroups previously thought to be predominantly symbiotic. Six FL*nif* strains from forest soil formed a clade within the *B. elkanii* supergroup, sister to known nodulating members (**Fig. S3**). Fourteen isolates were positioned at the basal nodes of the *B. japonicum* supergroup, representing the earliest-diverging FL*nif* lineage within this group (**Fig. S3**).

**Figure 1.**
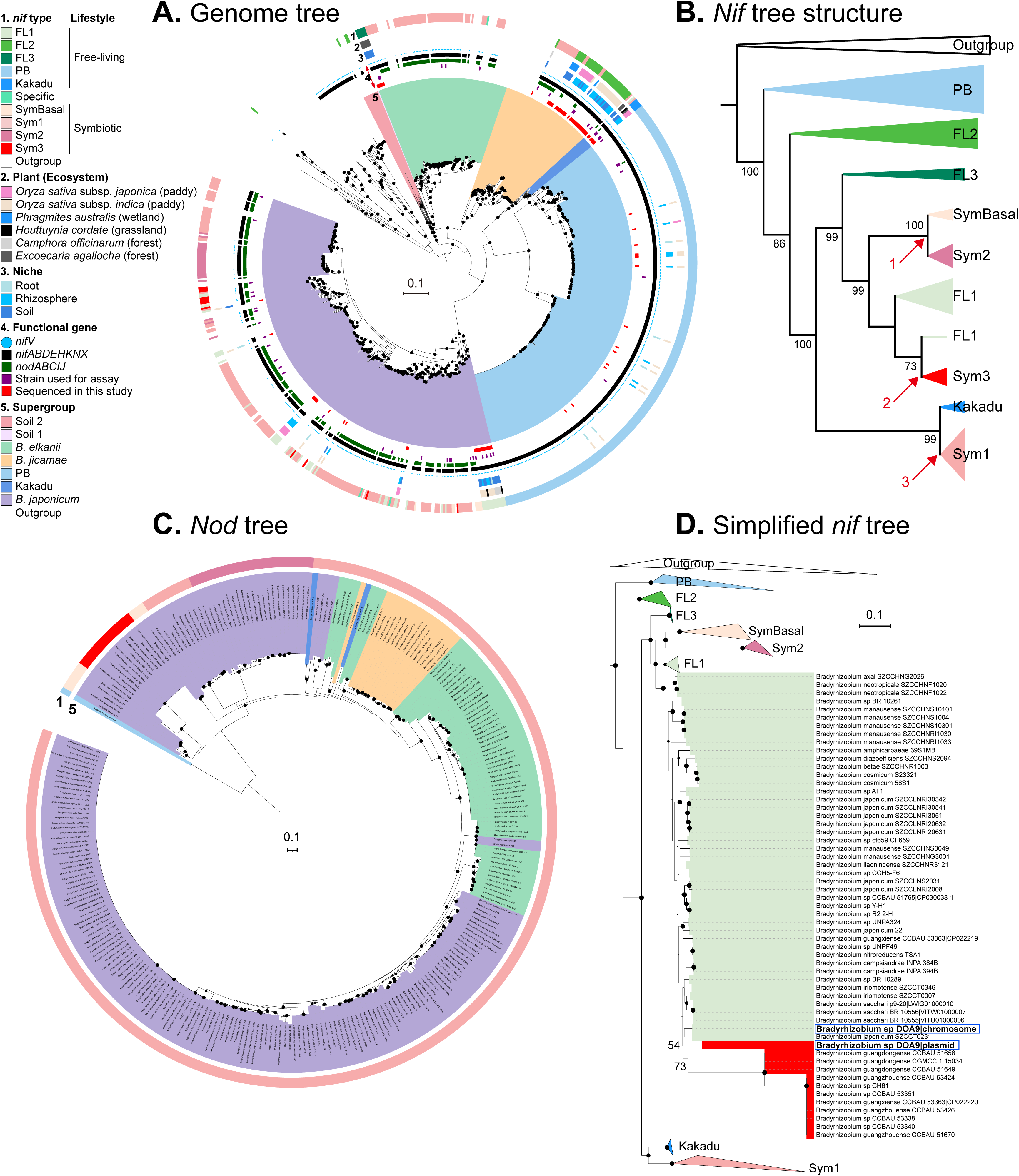
Phylogeny of *Bradyrhizobium* and evolutionary patterns of *nif* and *nod* genes. (A) Maximum-likelihood phylogenomic tree of *Bradyrhizobium* based on 123 orthologous genes identified in a previous study (22). Strains from Xanthobacteraceae were used as the outgroup. Members of the FL1, FL2, FL3, Kakadu, PB, SymBasal, Sym1, Sym2, and Sym3 clusters (refer to **Fig. S4**) are also labeled on this tree. (B) Concise *nif* gene tree illustrating the evolutionary relationships among different types of *nif* genes in *Bradyrhizobium*. The topology is derived from **Fig. S4**, with branch lengths omitted for clarity. Red arrows indicate lifestyle transitions from free-living to symbiosis. The ultrafast bootstrap values are shown for key nodes. (C) *Nod* phylogeny of *Bradyrhizobium* based on concatenated alignments of *nodABCIJ* under the LG+PMSF(C60)+G+F model, rooted with the minimum variance (MV) method. (D) Simplified *nif* phylogeny of *Bradyrhizobium* derived from **Fig. S4**, focusing on the evolutionary transition from the *B.* sp. DOA9 FL1 *nif* type to DOA9LSym3 *nif* type and to other Sym3 lineages. Other *nif* types are omitted for clarity. Black circles on nodes indicate ultrafast bootstrap values ≥ 95%, as calculated by IQ-Tree. The scale bar represents 0.1 nucleotide substitutions per site.

Another 35 FL*nif* members, isolated from forest, wetland, and paddy soils, were sporadically distributed within the *B. jicamae* supergroup **(Fig. 1A, S4)**. We also expanded the Kakadu supergroup with seven FL*nif* members from subtropical paddy and grassland, whereas only two symbiotic members from tropical legumes were previously known **(Fig. 1A, S4)** (34, 35).

By integrating these 88 new genomes with 710 publicly available *Bradyrhizobium* genomes and 96 outgroup members, we assembled a comprehensive dataset of 894 isolates (**Fig. S5, Dataset S1, S2**). This phylogenetic framework allowed reconstruction of the evolutionary history and functional adaptation of *nif* genes within the genus.

### Phylogenetic evidence for multiple independent origins of symbiotic nif clusters

To trace the evolution of the nitrogen-fixing trait, we constructed a phylogeny using concatenated *nif* genes (*nifABDEHKNX*) with the best-fitting substitution model LG+C60+G+F. This analysis assigned *nif*-carrying strains to nine phylogenetically distinct clusters (FL1, FL2, FL3, PB, Kakadu, SymBasal, Sym1, Sym2, and Sym3). One type (classified as “Specific” in **Fig. S4**), represented by three strains, is nested within a symbiotic clade (**Fig. S4, S6**). These strains lack detectable *nod* genes in their current genome assemblies, which may reflect incomplete genome assembly. Five clades (FL1, FL2, FL3, PB, and Kakadu) are capable of free-living nitrogen fixation, whereas the others are symbiotic (**Fig. 1B, S4**). Eight strains carried two *nif* clusters. Two of these strains carried two Sym1 clusters. The remaining six carried clusters belonging to different *nif* types; these are defined as dualL*nif* strains. Specifically, four carried one Sym1 and one FL1 cluster, and two carried one Sym3 and one FL1 cluster (**Dataset S2**). Our new isolates expanded FL1 (previously identified as FL cluster) and FL2 (previously containing only one strain, SZCCT0283, and not classified as a *nif* type) (22), and defined two new groups: FL3 and Kakadu (**Fig. 1B, S2, S4**).

The phylogenetic topology showed that the PB clade formed the earliest branching lineage, followed by the FL2 clade (**Fig. 1B, S4**). All other *nif* clades descended from a common ancestor shared with FL2 (**Fig. 1B**). The NF-dependent symbiotic *nif* lineages (SymBasal, Sym1, Sym2, and Sym3) were not monophyletic; they were interspersed among free-living clades. The *nod* gene phylogeny of *Bradyrhizobium* (**Fig. 1C, S7**) was incongruent with the *nif* tree (**Fig. 1B, S4**). Strains carrying Sym1 *nif* clusters formed a monophyletic group in the *nif* tree, but they were paraphyletic in the *nod* tree, where strains carrying Sym2 *nif* clusters were embedded within the Sym1 group. Strains that carried SymBasal and Sym3 *nif* clusters had *nod* genes that shared a recent common ancestor in the *nod* tree (**Fig. 1C**), yet these same strains were placed in separate subclades in the *nif* tree (**Fig. 1B**). The broader phylogenetic tree including other rhizobial genera (**Fig. S8**) showed that the symbiotic *Bradyrhizobium nif* clusters formed distinct, genus-specific lineages. These independently evolved symbiotic *nif* lineages did not show strict host specificity. Different symbiotic *nif* types co occurred within the same legume host genus, and the same *nif* type was found across multiple host genera (**Fig. S9**).

Among the six dualL*nif* strains, the two carrying Sym3 and FL1 clusters (*B*. sp. DOA9 and *B. guangxiense* CCBAU 53363) were particularly informative. In the *nif* gene tree (**Fig. 1D**), the Sym3 cluster of DOA9 formed the basal lineage within the Sym3 clade and was sister to a clade consisting of FL1 clusters from DOA9 and strain SZCCT0231. For the four strains carrying Sym1 and FL1 clusters, the Sym1 clusters were identical in phylogenetic position to typical Sym1 clusters (**Fig. S10**).

### Conserved architecture of free-living *nif* islands contrasts with variable symbiotic *nif* gene neighborhoods

Lineages capable of free-living nitrogen fixation (FL1, FL2, FL3, PB, Kakadu) shared a highly conserved *nif* island architecture despite their phylogenetic divergence (**Fig. 2A, S11**). The region was invariably flanked by *XDH* (encoding xanthine dehydrogenase) and *modD* (involved in molybdate transport) and maintained a consistent core gene order (**Fig. 2A, S11**): *nifA* (transcriptional activator) (36, 37), the *suf* cluster (Fe-S assembly) (38, 39), the structural nitrogenase genes *nifDKENX*, *nifHQ*, and *nifV* (40), the electron transfer genes *fixABCX* (41), and the molybdate transport *mod* cluster (42). The island also contained stress tolerance genes such as *glbO* (truncated hemoglobin for oxygen detoxification) (22, 43), *hspQ* (heat and oxidative stress resistance) (44, 45), and, in some clades, *TST* (cyanide detoxification) (46) and *MAO* (hydrogen peroxide generation) (47). Each free living lineage exhibited unique features: FL2 carried additional stress tolerance genes (*TST*, *MAO*, *hspQ*), whereas FL3 and Kakadu lacked these genes; *nifH* copy number and molybdate transporter composition (*modABCD* vs. *modCD*) also varied (**Fig. 2A, S11**).

**Figure 2.**
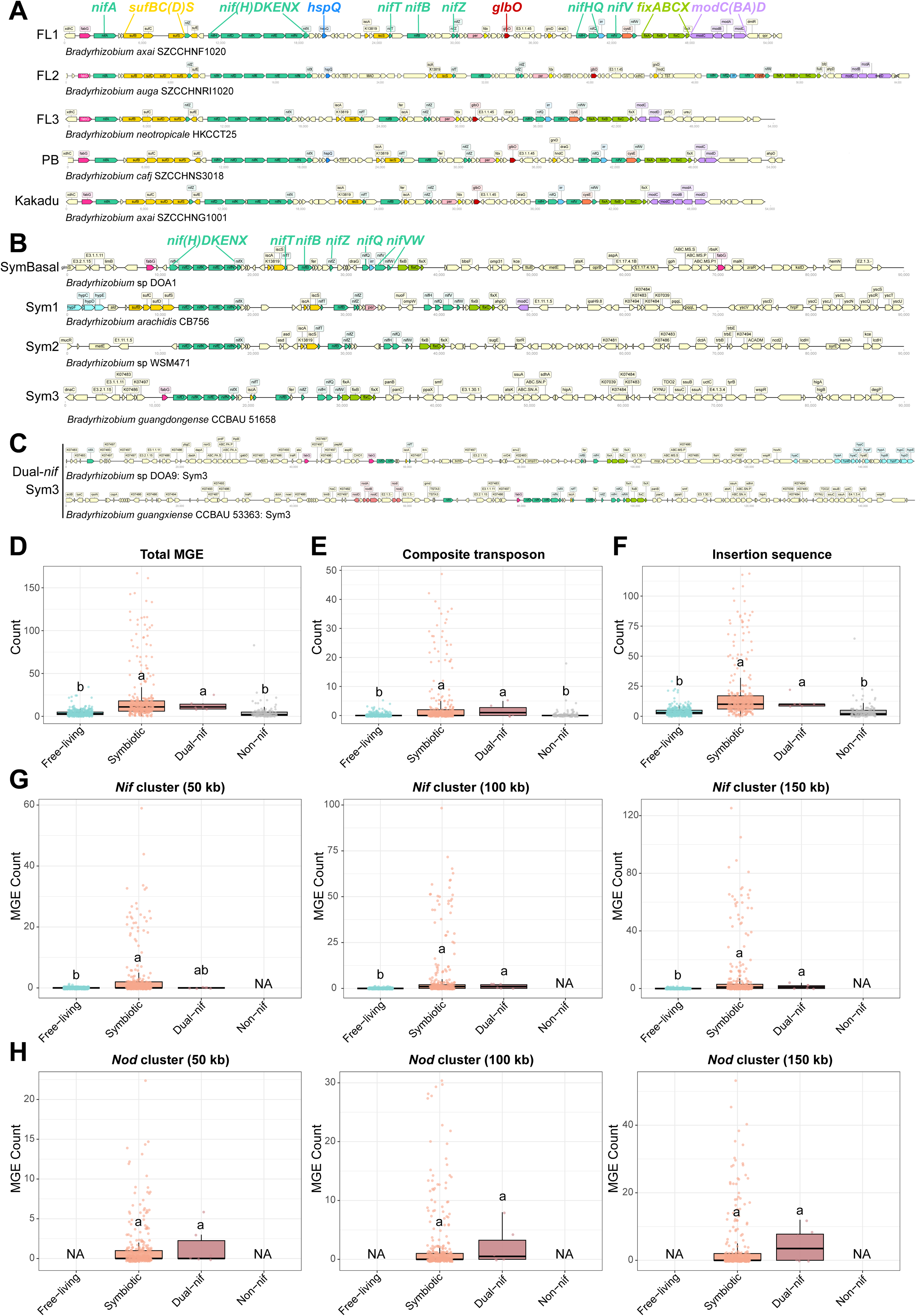
Genomic organization of *nif* clusters and mobile genetic elements (MGEs) abundance across lifestyles. (A-C) Gene arrangements of *nif* gene clusters in representative strains of three groups: (A) free living *nif* islands (FL*nif* strains: FL1, FL2, FL3, PB, Kakadu), (B) symbiosis islands (symbiotic strains: SymBasal, Sym1, Sym2, Sym3), and (C) Sym3 type of *nif* clusters from dual-*nif* strains. Genes with different functions are distinguished by different colors. Visualization was generated with dna-features-viewer v3.0.3 (81). In symbiosis island of symbiotic *Bradyrhizobium*, *nod* and T3SS genes are often located far from *nif* genes (9, 82) and are therefore not shown. “SymBasal” denotes the basal lineages of the nodulating *B. japonicum* supergroup (refer to Fig. 1A**, S3**). “Sym1, Sym2, and Sym3” represent different clusters of symbiotic *Bradyrhizobium* as indicated in Fig. 1B. (D–F) Number of MGEs across *Bradyrhizobium* with different lifestyles: (D) total MGEs, (E) composite transposons, and (F) insertion sequences. Each point represents one strain. Five outlier strains (*B.* sp. 1S5, 323S2, 38S5, 144S4, and *B. diazoefficiens* NK6) with exceptionally high counts are excluded from panels D-F for clarity but are included in statistical analysis. (G-H) Number of total MGEs within 50 kb, 100 kb, and 150 kb of the (G) *nif* and (H) *nod* gene clusters. Statistical significance among lifestyles was assessed using the non-parametric Kruskal-Wallis H test, followed by post-hoc pairwise Mann-Whitney U tests. *P*-values were adjusted for multiple comparisons and unbalanced group sizes using the Benjamini-Hochberg (FDR) method. Different letters denote significant differences between groups (*P* < 0.05). Complete MGE counts for each strain are provided in **Dataset S10**.

The *nif*-associated genomic regions of NF-dependent symbiotic lineages exhibited profound structural plasticity (**Fig. 2B, S12**). Gene order was highly variable and shuffled, and the boundaries of these regions were often associated with mobile genetic elements (MGEs) such as integrases (13, 14). A systematic analysis of MGEs across all genomes distinguished composite transposons and insertion sequences. Symbiotic and dualL*nif* strains harbored significantly more such MGEs than free living lineages, both genome wide (KruskalLWallis H test, *P* < 0.05; **Fig. 2D-F**) and within 50L150 kb flanking regions of *nif* and *nod* gene clusters (**Fig. 2G, H**). The *nif* cluster regions of all symbiotic strains also showed lower GC content relative to the genome average (mean ΔGC = −2.1% to −4.5%), whereas free living strains showed no such deviation (mean ΔGC = 0% to +0.5%; **Fig. S13**).

A consistent feature distinguishing all NFLdependent symbiotic clusters from free living counterparts was the universal loss of the *glbO* gene from the *nif* island (**Fig. 2B**). Although *nifV* (encoding homocitrate synthase) is frequently lost in rhizobia (48, 49), more than 60% of sequenced symbiotic *Bradyrhizobium* retain *nifV* (**Dataset S2**). In the two Sym3+FL1 strains, the FL1 cluster retained the compact free living architecture (**Fig. 2A, S14A**), whereas the Sym3 cluster followed either a divergent arrangement (*B*. sp. DOA9: *fabG*-*nifDK*-insertion of other genes-*nifB*-*nifH*-*fixABCX*) or the canonical symbiotic structure (*B. guangxiense* CCBAU 53363: *fabG*-*nifDKENX*-*nifB*-*nifHQW*-*fixABCX*, spanning ∼25 kb) (**Fig. 2C, S12, S14A**). In the four Sym1+FL1 strains, the Sym1 cluster resembled typical symbiotic Sym1 arrangements (**Fig. 2B, S12, S14B**), whereas the FL1 cluster was indistinguishable from free living FL1 (**Fig. 2A, S11, S14B**). The two strains carrying two Sym1 clusters showed nearly identical copies (**Fig. S14C**).

### Functional hierarchy of nitrogen fixation efficiency under free-living conditions

In a minimal semi-solid medium (buffered nodulation medium, BNM), symbiotic strains (SymBasal, Sym1, Sym2, and Sym3) exhibited significantly lower nitrogenase activity than strains from lineages capable of free-living nitrogen fixation (FL1, FL2, FL3, PB, Kakadu, and dual-*nif* strains Sym1+FL1 and Sym3+FL1) (PhylANOVA, a phylogenetic ANOVA accounting for evolutionary dependence among species, *P* = 0.0033; **Fig. 3A**). NF-dependent symbiotic strains lacking *nifV* showed no detectable activity; those retaining it displayed minimal or no nitrogen fixation capability (**Fig. 3A**). Among strains capable of free-living nitrogen fixation and dual-*nif* strains, PB strains showed the highest nitrogenase activity, but differences within this group were not statistically significant (two-way ANOVA with Tukey’s HSD, *P* > 0.05; **Fig. 3A, Dataset S3**).

**Figure 3.**
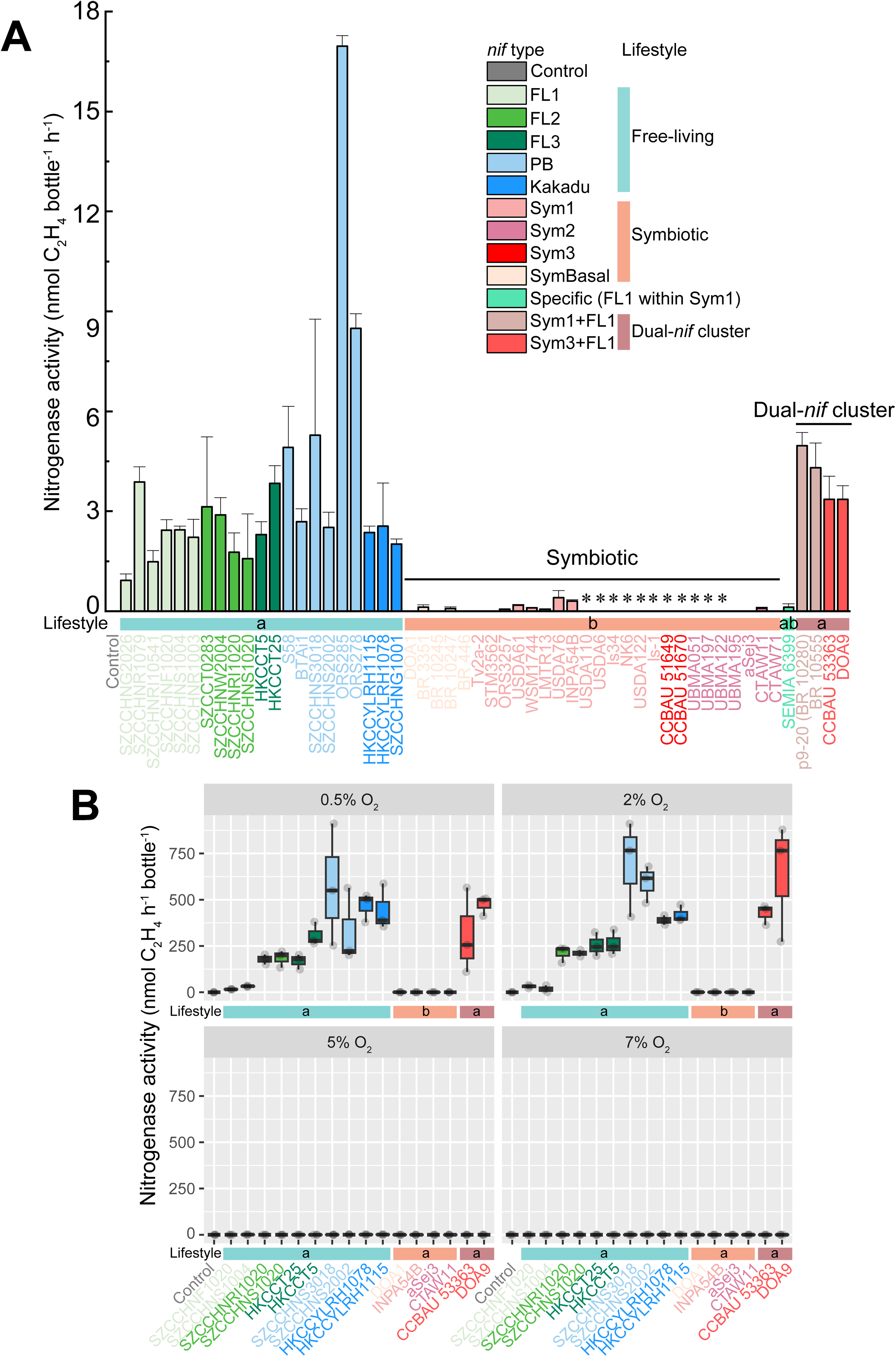
Nitrogenase activity of representative strains (3 biological replicates per strain) from diverse *nif* types *Bradyrhizobium* (FL1, FL2, FL3, PB, Kakadu, SymBasal, Sym1, Sym2, and Sym3) under free-living conditions measured by the acetylene reduction assay. (A) Several representative strains were grown for 6 days in 10 mL vacutainer tubes containing minimal growth N-free medium (BNM) with 10% acetylene under atmospheric air conditions. The column marked with an asterisk (*) indicates the strains lacking a *nifV* gene. (B) The representative strains were grown for 4 days in 65 mL vials containing N-free modified arabinose-gluconate (MAG) medium with 10% acetylene under varying O_2_ concentrations (0.5%, 2%, 5%, and 7%). Sym3 strains (except those possessing two *nif* clusters) were excluded in Fig. 3B due to the absence of *nifV*. The representative strains used in Fig. 3B from different *nif* types are presented in the following: SZCCHNF1020 (FL1), SZCCHNS1004 (FL1), SZCCHNS1020 (FL2), SZCCHNRI1020 (FL2), HKCCT5 (FL3), HKCCT25 (FL3), SZCCHNS3018 (PB), SZCCHNS2002 (PB), HKCCYLRH1078 (Kakadu), HKCCYLRH1115 (Kakadu), *B.* sp DOA1 (SymBasal), *B. forestalis* INPA54B (Sym1), *B. cytisi* CTAW11 (Sym2), *B. hipponense* aSej3 (Sym2), *B. guangxiense* CCBAU 53363 (dual-*nif* cluster, Sym3+FL1), and *B.* sp DOA9 (dual-*nif* cluster, Sym3+FL1). Significant differences among different lifestyles and *nif* type strains were determined by two-way ANOVA followed by Tukey’s test (*P* < 0.05). Error bars represent the standard error of the mean (n = 3). Different letters indicate statistically significant differences for each *nif* cluster were listed in **Dataset S3** and **S4** (*P* < 0.05).

To compare oxygen tolerance across divergent *nif* cluster architectures, we measured nitrogenase activity in liquid nitrogen-free modified arabinose-gluconate (MAG) medium under a controlled oxygen gradient (0.5%, 2%, 5%, 7% O_2_). All strains exhibited negligible nitrogenase activity at O_2_ concentrations of 5% or higher (**Fig. 3B, S15A, B**). At low oxygen (0.5% and 2% O_2_), a functional divide emerged. Strains from free-living *nif* clusters demonstrated significantly higher nitrogenase activity than NF-dependent symbiotic lineages (PhylANOVA, *P* = 0.03 and *P* = 0.04 in 0.5% and 2% O_2_ conditions, respectively; **Fig. 3B**). NF-dependent symbiotic strains showed virtually no activity (< 0.3 nmol C_2_H_4_ h^-1^ bottle^-1^) outside host nodules.

Although there was no statistically significant difference in nitrogenase activity among free-living clusters (PB, Kakadu, FL3, and FL2) under low oxygen (two-way ANOVA, Tukey HSD post-hoc test, *P* > 0.05; **Fig. 3B**, **Dataset S4**), they still exhibited a gradient of activity. PB strains exhibited the highest nitrogenase activity, with the Kakadu cluster performing comparably. FL3 and FL2 showed intermediate activity, reaching approximately 50% and 40% of the PB rate, respectively (**Fig. 3B**). The FL1 cluster showed nitrogenase activity significantly lower than all other free-living clusters (*P* < 0.05; **Fig. 3B, Dataset S4**) but remained above symbiotic strains (*P* < 0.005, **Fig. 3B, Dataset S4**).

### The oxygen-protective gene *glbO* in the *nif* island contributes to free-living nitrogen fixation

The *glbO* gene was consistently present in the *nif* island of lineages capable of free-living nitrogen fixation but universally absent from the symbiotic *nif* regions of NF-dependent symbiotic strains (**Fig. 2A-C, S4, S11, S12**). We tested whether *glbO* contributes to nitrogen fixation outside nodules using loss-of-function and gain-of-function approaches.

An insertional mutant of *glbO* was generated in the PB strain ORS285, which fixes nitrogen both free living and in symbiosis and uniquely carries *nod* genes (48, 50). In both semi-solid and liquid BNM media, the mutant ORS285Ω*glbO* showed significantly lower nitrogenase activity than the wild-type strain (one-way ANOVA with Tukey’s HSD, *P* < 0.005; **Fig. 4A, B**, **Dataset S5**). Ethylene formation kinetics were also drastically reduced in the mutant (**Fig. 4C, D**).

**Figure 4.**
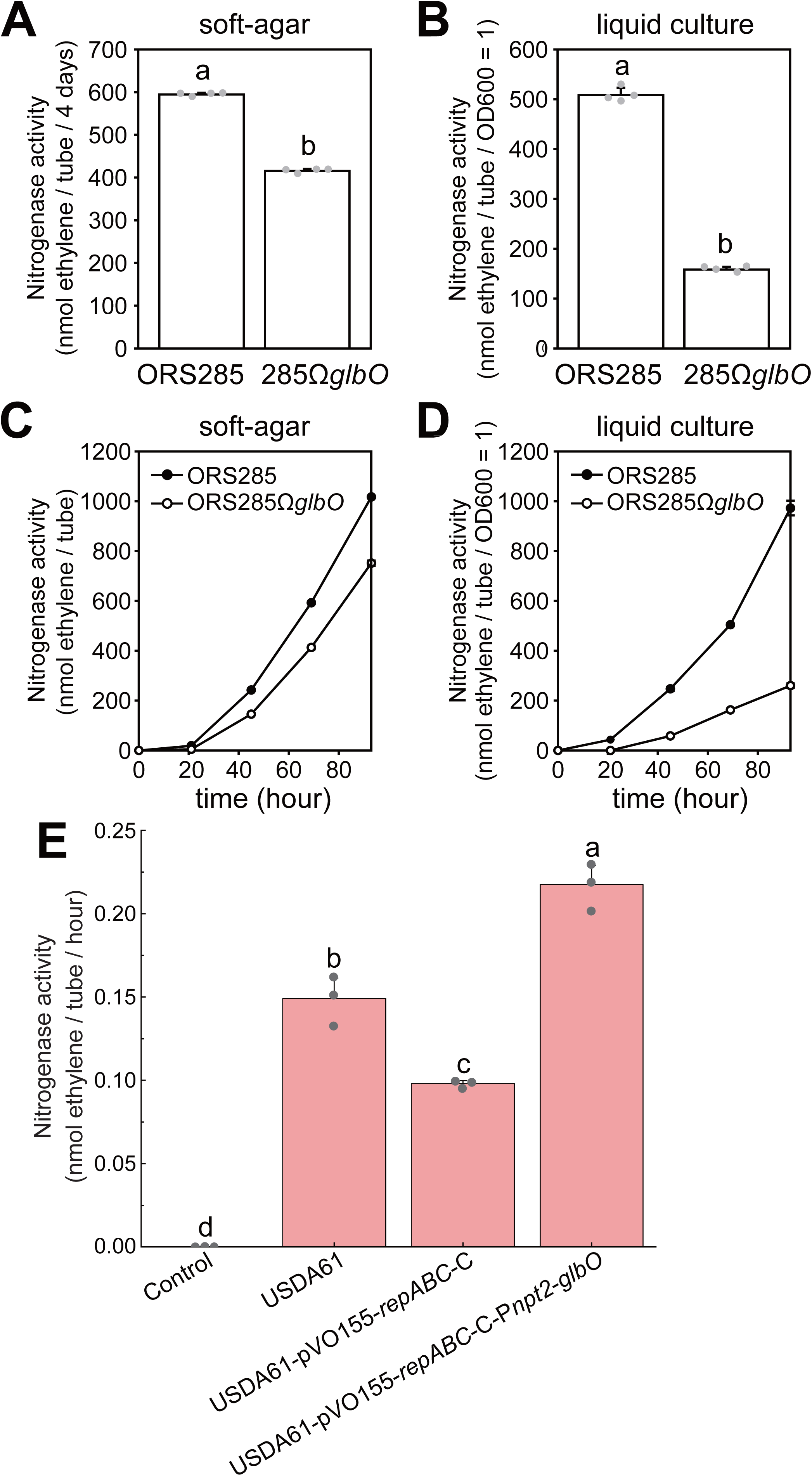
Nitrogenase activity of *Bradyrhizobium* wild-type and constructed strains assessed by acetylene reduction assay. Nitrogenase activity of the ORS285 and ORS285Ω*glbO* strains after 3 days of growth in BNM (A) soft-agar and (B) liquid culture (n = 4 biological replicates). The kinetics of ethylene formation by ORS285 and ORS285Ω*glbO* strain in BNM (C) soft-agar and (D) liquid culture (n = 2). (E) Nitrogenase activity of wild-type strain USDA61 and its complemented derivatives carrying the *glbO* gene expressed from a constitutive promoter (*npt2*) using the vector backbone pVO155-P*npt2-bjGFP*-*repABC*-C-NK6. Error bars represent the standard deviation. Different letters above the bars indicate statistically significant differences between the wild-type and corresponding constructed strains (**Dataset S5** and **S6**). Significant differences in Fig. 4A, 4B, and **4E** were determined by one-way ANOVA followed by Tukey’s post-hoc test (*P* < 0.0005).

The systematic absence of the *glbO* gene in NF-dependent symbiotic lineages suggests this gene is not essential for nitrogen fixation during symbiosis. To test this hypothesis, ORS285 and ORS285Ω*glbO* were inoculated into two *Aeschynomene* species: *A. indica* (which uses a NFLindependent pathway, **Fig. 5**) and *A. afraspera* (which uses an NFLdependent pathway, **Fig. S16**) (8). In both plants, the mutant and wild type strains showed no significant difference in host plant growth, nodule number, or plant nitrogenase activity (**Fig. 5ADC**, **S16A**D**C**). Microscopy confirmed that nodules induced by both strains were fully infected, with bacteria differentiated into bacteroids, the active nitrogen-fixing form during symbiosis (**Fig. 5D-K, S16D-K**).

**Figure 5.**
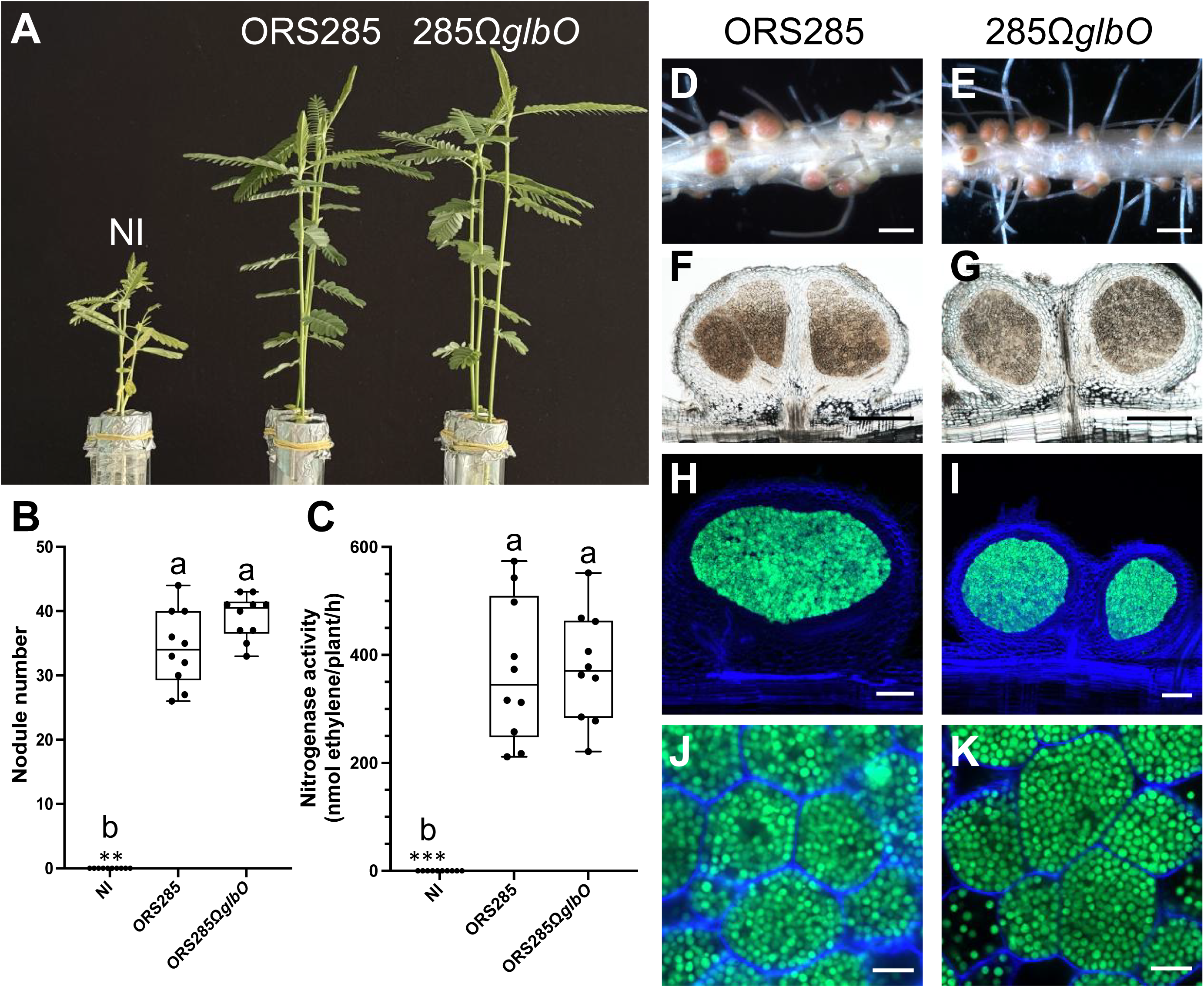
The symbiotic properties of the ORS285 and its *glbO* derivative mutant ORS285Ω*glbO* strains on *A. indica*. (A) Photo of *A. indica* aerial parts at 14 days post-inoculation (dpi) with ORS285 and ORS285Ω*glbO* strains. (B) Nodule number and (C) nitrogenase activity of *A. indica* plants at 14 dpi inoculated with ORS285 and ORS285Ω*glbO* strains. NI: non-inoculated. These box plots show the results obtained on 10 plants. Significant differences observed in Fig. 5B and Fig. 5C were tested using a non-parametric Krustal-Wallis test. Different letters above the bars indicate statistically significant differences. ** *P* < 0.005, *** *P* < 0.0005. Photo of roots and nodules induced by (D) ORS285 and (E) ORS285Ω*glbO* strains. Scale bars: 0.2 cm. Cross section of nodules induced by ORS285 (F) and (G) ORS285Ω*glbO* strains observed using light microscopy showing the internal infected tissue and the peripheral vascularization. Scale bars: 500 μm. Confocal microscopy images of sections of nodules induced by ORS285 (H, J) and ORS285Ω*glbO* (I, K) strains. The strains were tagged with GFP and calcofluor staining was used to observe plant cell walls (blue, plant cell wall). Scale bars: H and I, 200 µm; J and K, 10 µm.

To test whether introducing *glbO* into a symbiotic strain could restore free living nitrogenase activity, we expressed the *glbO* gene from ORS278 (a free living PB strain) under a constitutive npt2 promoter in *B. elkanii* USDA61, which naturally lacked *glbO* in its symbiotic island but carried *nifV*. The introduced plasmid was stable maintenance in USDA61 (**Fig. S17**). The transformed USDA61 strain showed a significant increase in nitrogenase activity under free-living conditions compared to the wild type and empty-vector control (one-way ANOVA with Tukey’s HSD, *P* < 0.05; **Fig. 4E, Dataset S6**). The maximal activity achieved (0.20 nmol C_2_H_4_ h^-1^ bottle^-1^) was significantly higher than that of the wild-type strain (*P* < 0.05, 0.15 nmol C_2_H_4_ h^-1^ bottle^-1^) but remained markedly lower than that of strains naturally harboring *glbO* in the *nif* island (e.g., PB strains, 2.5L17.0 nmol C_2_H_4_ h^-1^ bottle^-1^).

## Discussion

### Evolutionary history: free living nitrogen fixation as ancestral, symbiosis as derived and polyphyletic

Research on rhizobial evolution has long been dominated by a legume-centric paradigm that regard free-living nitrogen-fixing strains as derived outliers rather than central players (12–14). This study challenges that view. The *nif* gene phylogeny shows that the earliest-diverging clades (PB and FL2) are capable of free-living nitrogen fixation, establishing the *nif* island associated with this lifestyle as the ancestral state within *Bradyrhizobium*. This finding refutes the scenario of a single symbiotic ancestor for the genus (Scenario 1). The broader phylogeny including other rhizobial genera shows that symbiotic *Bradyrhizobium nif* clusters form distinct, genus-specific lineages, arguing against horizontal acquisition from other genera such as *Rhizobium*, *Sinorhizobium*, or *Mesorhizobium* (Scenario 4). The *nif* gene phylogeny further demonstrates that NF-dependent symbiotic lineages are polyphyletic, arising independently at least three times from distinct free-living ancestors. This pattern of multiple convergent origins is incompatible with a model of a single origin and subsequent radiation within the genus (Scenario 3).

DualL*nif* strains provide direct evidence for the transition from free living to symbiotic states. In *B*. sp. DOA9 (Sym3+FL1), the Sym3 cluster forms the basal lineage within the Sym3 clade and is sister to a clade consisting of FL1 clusters from DOA9 and strain SZCCT0231 (**Fig. 1D**). This phylogenetic position identifies DOA9 Sym3 as the transitional lineage from FL1 to Sym3. The gene arrangement of DOA9 Sym3 is highly divergent (*fabG*-*nifDK*-insertion of other genes-*nifB*-*nifH*-*fixABCX*) compared to the canonical Sym3 structure seen in other strains (e.g., *B. guangxiense* CCBAU 53363: *fabG*-*nifDKENX*-*nifB*-*nifHQW*-*fixABCX*) (**Fig. 2C, S10**). This unusual architecture, together with its basal phylogenetic position, supports the interpretation that DOA9 Sym3 represents an early stage in the evolution of a symbiotic *nif* cluster from a free living ancestor.

The *nod* gene phylogeny is incongruent with the *nif* tree (**Fig. 1B, C, S4**), arguing against co acquisition of *nif* and *nod* genes via a single symbiosis island. Strains carrying Sym1 *nif* clusters form a monophyletic group in the *nif* tree but are paraphyletic in the *nod* tree, with Sym2 strains embedded within Sym1. SymBasal and Sym3 *nif* type strains carry *nod* genes that share a recent common ancestor, yet these strains are placed in separate subclades in the *nif* tree. These patterns, together with the genomic evidence below, indicate that the capacity for free living nitrogen fixation is ancestral, and NFLdependent symbiotic competence is a derived trait that evolved convergently multiple times via independent acquisition of symbiosis islands by distinct free living ancestors (Scenario 2).

### Legume host selection: driver of transfer and maintenance but limited cross type exchange

Legume hosts exert strong selective pressure on symbiotic *Bradyrhizobium* (51, 52). Each independent transition from a free living ancestor to an NFLdependent symbiotic lifestyle (at least three origins documented by the *nif* phylogeny) may have been driven by legume host selection. After a new symbiotic *nif* type arises, host plants could facilitate its spread among compatible strains (53, 54).

Several lines of evidence are consistent with this interpretation. Symbiotic and dualL*nif* strains harbor significantly more mobile genetic elements (MGEs) than free□living lineages, both genome□wide and within flanking regions of *nif* and *nod* gene clusters (**Fig. 2D-H**). The *nif* cluster regions of symbiotic strains also show lower GC content relative to the genome average, whereas free living strains show no such deviation (**Fig. S13**). These signatures are consistent with horizontal acquisition of these regions as mobile genomic islands (55, 56). Gene order within the *nif*-associated region is highly conserved among strains carrying the same symbiotic *nif* type (SymBasal, Sym1, Sym2, or Sym3) but differs markedly between types (**Fig. 2B, S12**). This pattern indicates that each symbiotic *nif* type has evolved independently, with limited exchange of genetic material across different types. Further supporting this, many strains that are closely related in the phylogenomic tree carry different *nif* types (**Fig. 1A**), suggesting that horizontal transfer of *nif*Lassociated regions did not occur among such closely related strains.

The transitional DOA9 Sym3 cluster, described above, provides additional evidence for host driven architectural evolution. Its highly divergent gene arrangement differs from the canonical Sym3 structure found in other strains. This unusual arrangement likely represents an early stage after the transition from FL1 to Sym3. The subsequent evolution toward the canonical Sym3 arrangement observed in other Sym3 strains may have been favored by host selection for a more streamlined symbiotic *nif* cluster.

The distribution of *nif* types across host genera rules out strict co evolution or co-speciation between a particular *nif* type and a specific host lineage. Different symbiotic *nif* types co occur within the same host genus (e.g., Sym1 and Sym2 from *Lupinus*; Sym1 and Sym3 from *Arachis*; Sym1 and SymBasal from *Vigna*, *Stylosanthes*, *Centrolobium*, and *Aeschynomene*), and the same *nif* type (e.g., Sym1) can be found across multiple host genera (**Fig. S9, Dataset S2**). These patterns indicate that once a symbiotic *nif* type has arisen (potentially through host driven transition from a free living ancestor), it can subsequently spread to diverse legume hosts, and different *nif* types can colonize the same host. Thus, while host selection may drive each independent origin and subsequent refinement (e.g., DOA9 FL1 to DOA9 Sym3 to canonical Sym3), it does not impose a strict one to one association between a *nif* type and a host genus.

Taken together, the data support a model in which legume hosts are powerful drivers of the origin, dissemination, and architectural refinement of symbiotic *nif* clusters (51). The observation that distinct *nif* types maintain their integrity across diverse host genera, and that different types can co exist within the same host plant, suggests that horizontal exchange of symbiotic regions is not readily shared across different *nif* types.

### Functional, genomic, and molecular dichotomy between free living and symbiotic *nif* systems

Free living and symbiotic lineages differ profoundly in their ability to fix nitrogen outside the host, in the genomic architecture of their *nif* regions, and in their requirement for specific protective genes.

Strains capable of free living nitrogen fixation (including dualL*nif* strains) exhibit robust nitrogenase activity under free living conditions, whereas symbiotic strains show virtually no activity outside nodules (**Fig. 3A, B**). This confirms that symbiotic lineages have undergone evolutionary specialization to the nodule niche, losing the capacity for independent nitrogen fixation (57, 58). Among free living clusters, PB strains show the highest activity, with Kakadu, FL3, and FL2 achieving intermediate levels. The FL1 cluster retains genetic potential but shows constrained efficacy, consistent with its phylogenetic position as a transitional group from which Sym3 evolved.

Free living lineages share a highly conserved *nif* island architecture (*nifA*L*suf*L*nifDKENX*L*glbO*L*nifHQ*L*nifV*L*fixABCX*L*mod*), flanked by *XDH* and *modD* (**Fig. 2A, S11**). This conservation is consistent with vertical inheritance from a common ancestor. In contrast, symbiotic *nif*Lassociated regions are highly variable in gene order and composition (**Fig. 2B, S12**), often associated with MGEs (**Fig. 2D-H**), and have lower GC content (**Fig. S13**). This variability reflects independent capture, rearrangement, and streamlining of the ancestral *nif* island upon each transition to symbiosis. The process involved integration of novel symbiosis-promoting genes (*nod*, T3SS genes) via horizontal gene transfer (HGT), together with the loss of genes that became dispensable within the protected nodule niche (59). One such gene is *nifV*. Around 40% of sequenced NF-dependent symbiotic *Bradyrhizobium* strains lack *nifV* (**Fig. 2A, B**), yet its loss aligns with the known ability of legume hosts to supply homocitrate (48), a classic example of reductive evolution and metabolic integration.

The *glbO* gene shows a universal pattern across all *Bradyrhizobium* lineages. In every free living lineage (FL1, FL2, FL3, PB, Kakadu), *glbO* is located within the *nif* island, within 60 kb of the *nif* genes. In NF-dependent symbiotic strains, *glbO* is universally absent from the *nif*Lassociated region. Even when a *glbO* homolog is present elsewhere in the genome, it resides far from the *nif* genes (often > 200 kb away or on a different contig).

Loss of function mutagenesis of *glbO* in the PB strain ORS285 significantly reduced nitrogenase activity under free living conditions. Gain of function expression of *glbO* in the symbiotic strain USDA61 increased free-living nitrogenase activity, although not to native levels (**Fig. 4**). The same *glbO* mutant showed no defect in symbiosis with two *Aeschynomene* species, forming fully effective nodules (**Fig. 5, S14**). These results establish that *glbO* is required for optimal nitrogen fixation under fluctuating oxygen conditions outside nodules but is dispensable inside the protected, microaerobic nodule environment (60, 61). The failure of *glbO* introduction to fully restore free living activity in USDA61 suggests that symbiotic lineages have lost additional factors, possibly including other stress protection genes, that are essential for robust nitrogenase activity in the absence of the host (51). We cannot exclude the possibility that the relatively modest contribution of *glbO* is influenced by the use of a constitutive promoter, which may not fully recapitulate its native regulatory dynamics. An *in silico* analysis of the upstream region of *glbO* in ORS278 reveals putative NifA and RpoN binding motifs associated with the promoter region of *nifB*, located approximately 14 genes upstream. This genomic context is consistent with the possibility of coordinated regulation within a broader *nif* regulatory landscape.

### Evolutionary and functional divergence of *nif* clusters

The PB lineage differs from other *Bradyrhizobium* supergroups in its pattern of *nif* inheritance. The PB clade forms a monophyletic group in both the core genome tree and the *nif* gene tree, indicating a vertically inherited *nif* island within an ecologically coherent lineage. This stability may be linked to specialization in unique niches such as stem nodulation and waterlogged soils (62, 63), which could reduce genetic exchange with other soil microbiota. In other major supergroups (e.g., *B. japonicum*, *B. elkanii*), *nif* gene clusters are interspersed among lineages with different core genomes, revealing a mosaic pattern. This pattern reflects the role of HGT in distributing symbiotic competence across diverse genetic backgrounds. The PB lineage thus represents an evolutionarily more insulated group with a fixed ancestral *nif* apparatus, whereas in other supergroups, *nif* islands behave as MGEs.

PB strains are *nod*Lfree but nodulating, yet they have not undergone the same degree of reductive evolution seen in other symbiotic lineages. This difference correlates with their unique NFLindependent nodulation with *Aeschynomene*. PB strains form nodules on both roots and stems. Stem nodules are exposed to higher oxygen tensions due to atmospheric contact and oxygen production from chloroplast rich epidermal cells (64). This elevated oxygen environment likely favors the retention of free living nitrogen fixation capabilities. PB strains are also prevalent in rice paddy soils, where free living nitrogen fixation provides a selective advantage (23).

Among all lineages capable of free-living nitrogen fixation, PB and dual-*nif* strains exhibit the highest nitrogenase activity (**Fig. 3A, B**). The PB cluster carries additional stress-tolerance genes (*TST*, *MAO*, *hspQ*) and a potential duplicated *nifH* (**Fig. 2A, S11**). These features make PB strains promising candidates for plant growth promoting bacteria in waterlogged crops such as rice (23, 26, 27). Dual-*nif* strains (Sym1+FL1 and Sym3+FL1) carry increased core *nif* gene copies. In these strains, the symbiotic *nif* cluster operates inside nodules whereas the FL1 cluster remains active under free living conditions (21, 57, 65).

Their coexistence may allow a broader range of environmental responsiveness, although the underlying regulatory mechanisms remain to be tested. The FL1 cluster exhibits a variable phenotype. Its nitrogenase activity is robust in semi-solid BNM medium (which approximates nodule conditions) (48, 66) but significantly reduced in liquid MAG medium (which facilitates oxygen diffusion) (**Fig. 3A, B**). This pattern is consistent with the phylogenetic position of FL1 as a paraphyletic group from which the symbiotic Sym3 clade evolved (**Fig. 1B**). Thus, the FL1 cluster retains high nitrogen fixation capacity under conditions mimicking the symbiotic niche yet exhibits constrained efficacy under fully free-living conditions.

### Ecological and agricultural implications

Phylogenetic analyses expand the known ecological strategies of *Bradyrhizobium* by revealing basal free-living nitrogen-fixing members within the *B. japonicum* and *B. elkanii* supergroups (**Fig. 1A, S3**) previously thought to be predominantly symbiotic (10, 18, 67). This suggests a broader and potentially ancestral role for these members in environmental nitrogen cycling than previously appreciated (21–23). Given the global prevalence of *Bradyrhizobium* in soils and non-legume plants, the community of strains capable of free living nitrogen fixation is genetically and functionally rich, with significant potential contribution to nitrogen budgets beyond legume symbiosis.

The high nitrogen fixation efficiency of PB strains, together with their ability to colonize rice and other non legume crops endophytically (23, 25, 26), points to their immediate promise for sustainable agriculture. These strains could enhance nitrogen availability for non-legume crops through rhizosphere colonization or endophytic associations. This work establishes a new evolutionary paradigm, positioning free-living nitrogen-fixing *Bradyrhizobium* not as derived outliers but as the ancestral source from which symbiotic lineages repeatedly emerged. Harnessing these ancestral traits offers a promising strategy for developing novel biofertilizers to reduce dependence on synthetic nitrogen fertilizers.

## Materials and Methods

### Sample collection, bacterial isolation and genome sequencing

To substantially expand the genomic representation of FL*nif Bradyrhizobium*, we collected root and soil samples from a diverse array of non-legume plants across major ecosystems. Our sampling strategy targeted five geographically distinct regions in China with varying climatic conditions to capture broad ecological diversity (**Fig. S1**). From croplands, we sampled rice (*Oryza sativa* subsp. *indica* and *japonica*) and maize (*Zea mays*); from grassland, forest, and wetland ecosystems, we collected *Houttuynia cordata*, *Camphora officinarum*, *Excoecaria agallocha*, and *Phragmites australis*. However, *Bradyrhizobium* was not isolated from the maize samples collected in the Shanxi and Anhui provinces (**Fig. S1, Dataset S1**).

For each sample, we processed three distinct niches: the root endosphere, rhizosphere, and bulk soil. Bacterial isolation was performed on selective Eosin-Methylene blue Modified Arabinose-Gluconate (EM-MAG) medium (22), yielding 88 novel *Bradyrhizobium* isolates. We characterized basic soil properties, including pH, nutrient content, and heavy metal availability, for all sample sites to provide ecological context (**Dataset S1**). Genomic DNA from each isolate was extracted and sequenced using the MGISEQ-2000 PE150+150+10+10 platform. Raw reads were processed and assembled into high-quality draft genomes as described previously (23). The quality of genome assemblies (e.g., completeness and contamination) (**Dataset S7**) was assessed by CheckM v1.0.7 with default parameters (68).

### Phylogenomic and evolutionary analysis

We constructed a comprehensive phylogenomic tree of *Bradyrhizobium* (**Fig. 1A**) to elucidate the diversity and distribution of *Bradyrhizobium*. Our dataset incorporated the 88 newly sequenced genomes with 806 publicly available genomes from *Bradyrhizobium* and related outgroups, totaling 894 isolates (**Fig. S5, Datasets S1, S2**). This phylogeny was inferred from 123 shared single-copy genes identified in our previous *Bradyrhizobium* phylogenomic study (22) using IQ-Tree v2.2.0 (69).

To specifically trace the evolution of the *nif* genes, we built a phylogeny (**Fig. S4, S6**) based on the concatenated alignments of the amino acid sequences of eight core *nif* genes (*nifABDEHKNX*). We employed a rigorous model-testing approach, comparing four complex substitution models (LG+G+F, LG+PMSF(C20)+G+F, LG+PMSF(C40)+G+F, LG+PMSF(C60)+G+F) and selected the best-fitting model (LG+C60+G+F) based on AIC (Akaike information criterion) or BIC (Bayesian information criterion) scores (**Dataset S8**). PMSF indicates the posterior mean site frequency approximation for mixture substitution models (70, 71). To investigate the evolution of the *nod* genes, *nod* gene phylogeny was constructed from concatenated alignments of *nodABC* and *nodABCIJ* using the following substitution models: LG+G+F, LG+C20+G+F, and LG+C60+G+F. All trees were visualized and edited in iTOL (https://itol.embl.de) (72). Mobile genetic elements (MGEs) were identified using MGEfinder (version 1.0.6) across all Bradyrhizobium genomes (73).

*Bradyrhizobium* strains were classified based on the presence of *nif* (*nifABDEHKNX*) and *nod* (*nodABCIJ*) genes into NF-dependent symbiotic (both, allowing a maximum of two missing genes), *nif*-carrying strains capable of free-living nitrogen fixation (FL*nif*, only *nif*), and free-living non-*nif* carrying (FLnon*nif*, neither) categories (**Fig. 1A, S5, Dataset S1, S2**). Specifically, there are some special FL*nif* members that are *nod*-free but nodulating members (traditionally called Photosynthetic *Bradyrhizobium* supergroup). These members lack *nod* genes but can form nodules with *Aeschynomene* (23, 50).

### Genetic constructs and bacterial strain engineering

To functionally investigate the role of the *glbO* gene, we employed both loss-of-function and gain-of-function approaches. An insertional mutant of *glbO* in *B.* sp. ORS285 (ORS285Ω*glbO*) was generated via single-crossover homologous recombination using the non-replicative pVO155*-Sm-*P*npt2-GFP* vector harboring an internal *glbO* fragment (65).

For the introduction of the *glbO* gene into the NF-dependent symbiotic strain *Bradyrhizobium elkanii* USDA61, we followed a strategy inspired by a previous study (74) showing stable replication of plasmids carrying *repABC* genes in diverse rhizobia. For this purpose, the *repABC* region from an indigenous plasmid of *B. diazoefficiens* NK6 (75) was cloned into the non-replicative plasmid pVO155-P*npt2-bjGFP* harboring a kanamycin resistant gene and a constitutive *GFP* which codon composition was optimized for *Bradyrhizobium* (9, 76). The *glbO* gene from *B.* sp. ORS278 strain under the constitutive *npt2* promoter was then cloned into pVO155-P*npt2-bjGFP*-*repABC*-C-NK6 and introduced into USDA61 by electroporation. All plasmids, primers, and cloning strategies are detailed in Supplementary Text and **Dataset S9**.

### Physiological assays

To systematically assess and compare the nitrogenase activity across the distinct *nif* clusters identified in our study, we used the acetylene reduction assay (ARA) to evaluate general capacity and oxygen sensitivity. We initially profiled several representative strains from each *nif* cluster in a semi-solid minimal growth medium (buffered nodulation medium [BNM], which is a synthetic plant growth medium) (48, 66, 77) under non-oxygen-controlled conditions to determine a baseline for free-living nitrogenase activity (**Fig. 3A**). To precisely delineate the oxygen tolerance intrinsic to each cluster’s genomic architecture, a more refined assay was conducted in a nitrogen-free MAG liquid medium (22, 78) under a controlled gradient of oxygen tensions (0.5%, 2%, 5%, and 7% O_2_), thereby defining the functional window for nitrogen fixation in a free-living context (**Fig. 3B**). The functional characterization of the *glbO* insertional mutant (ORS285Ω*glbO*) and its wild-type progenitor encompassed both ARA and kinetic studies, executed in both semi-solid and liquid BNM media under non-oxygen-controlled conditions (**Fig. 4A-D**). To evaluate the gain-of-function, we measured the nitrogenase activity in USDA61 and its mutant in a semi-solid BNM medium (**Fig. 4E**). To verify the role of *glbO* in a symbiotic condition, plant inoculation experiments were conducted with both *A. indica* and *A. afraspera*.

### Statistical analysis

The non-parametric Kruskal-Wallis H test, followed by post-hoc pairwise Mann-Whitney U tests, were used to assess the differences in MGE counts among lifestyles and *nif* types (**Fig. 2C-G**). We performed a phylogenetic ANOVA (phylANOVA) analysis (79) to compare the differences in nitrogenase activity among *Bradyrhizobium* with different phenotypic traits (i.e., capable of free-living nitrogen fixation vs. symbiotic nitrogen fixation) (**Fig. 3A, B**).

Since nitrogenase activity was measured by acetylene reduction assays under free-living conditions and dual-*nif* cluster strains exhibited higher nitrogenase activity under these conditions, they were classified as strains capable of free-living nitrogen fixation in phylANOVA analysis. To resolve differences in nitrogenase activity among various lifestyles and *nif* types of *Bradyrhizobium*, a linear mixed-effects model (LMM) was applied to account for two fixed effects (lifestyle and *nif* type) and two random effects (strain identity and biological replication), ensuring robust inference of significant differences (80) (**Fig. 3A, B**).

## Data Availability

The genomic sequences and raw reads of the 88 newly sequenced *Bradyrhizobium* isolates (i.e., missing 2 raw reads for 1 genome: HKCCT186) are available under the NCBI BioProject ID: PRJNA1450941 (https://dataview.ncbi.nlm.nih.gov/object/PRJNA1450941?reviewer=7ggij0q9hn2cev2fbgf4ojif3o), while the assembled sequence of the missed genome is available at NCBI BioProject ID: PRJNA1395771 and will be released in the next version of the manuscript.

## Acknowledgements

We thank Xiaoyuan Feng for assembling the genomes and Xingqin Lin for discussing the assay design. H.L. was supported by the Hong Kong Research Grants Council Area of Excellence Scheme (AoE/M-403/16) and the Hong Kong Research Grants Council General Research Fund (14107823); E.G. was supported by a grant from the French National Research Agency (“ET-Nod”; ANR-20-CE20-0012).

