## Supplementary material for "The free-living wellspring of symbiotic nitrogen fixation in *Bradyrhizobium*": SI figures

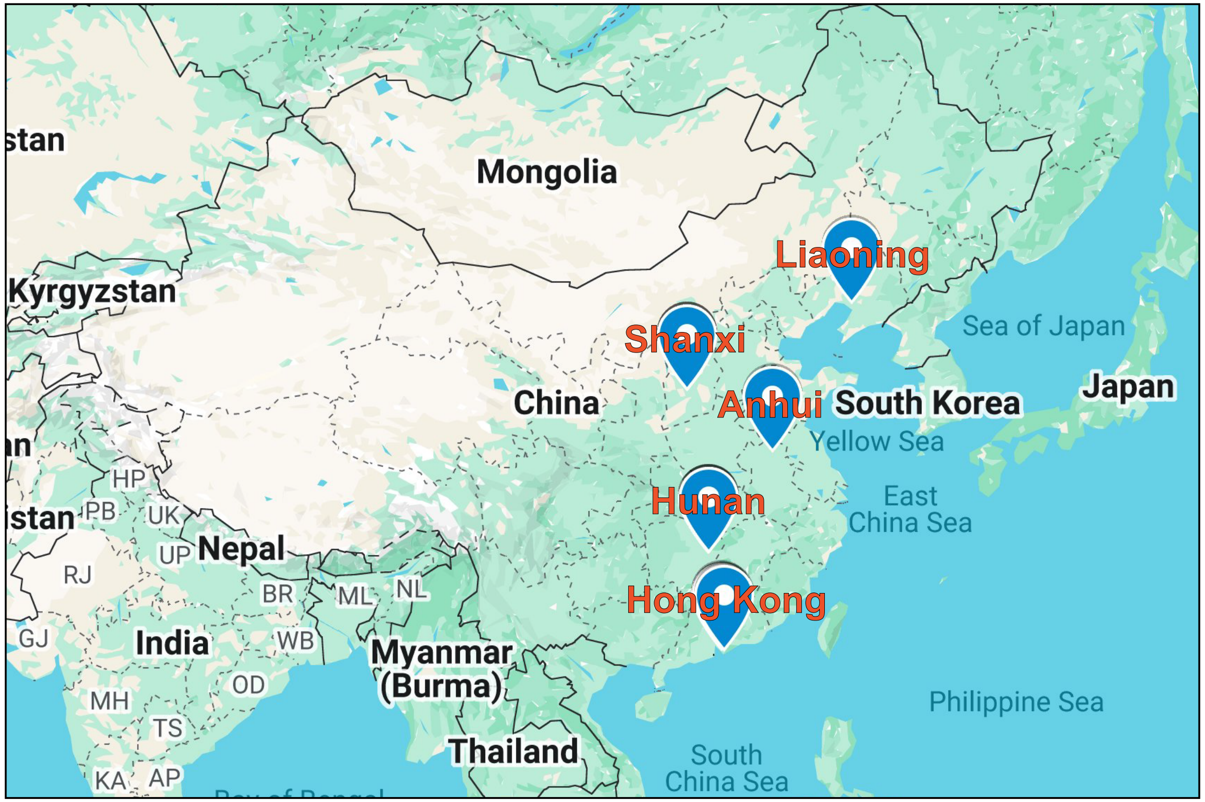


**Figure S1.** Geographic distribution of five sampling sites where seven plant species and nearby soils were collected. Note that rice (*Oryza sativa* subsp. *indica*) samples were obtained from Hunan (27.948 °N, 113.221 °E) province, while another rice (*Oryza sativa* subsp. *japonica*) samples were collected from Liaoning (40.826 °N, 122.209 °E) province and Hong Kong (22.418 °N, 114.080 °E). Maize (*Zea mays*) samples were sampled from Anhui (33.385 °N, 117.255 °E) and Shanxi (36.568 °N, 111.876 °E) provinces. *Houttuynia cordata* and *Camphora officinarum* were gathered from Hunan grassland (27.943 °N, 113.225 °E) and forest (27.944 °N, 113.226 °E), respectively.

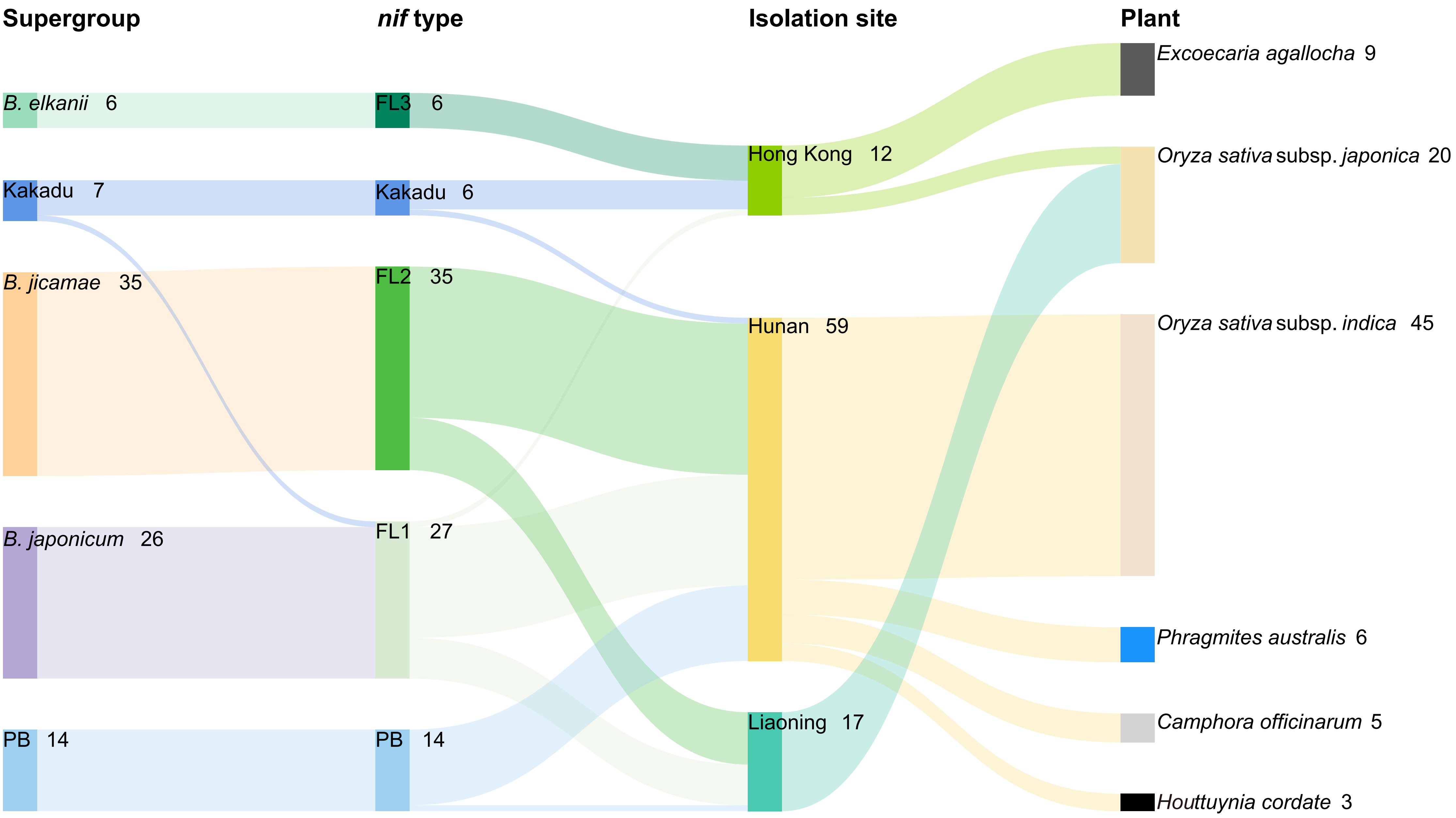


**Figure S2.** Distribution of *Bradyrhizobium* isolates across phylogenetic supergroups, *nif* types, isolation sites, and plant hosts. Sankey diagram illustrates the flow of isolate counts from left to right through four categorical levels: supergroup (as defined in **Fig. 1A**), *nif* type (as defined in **Fig. S4**), isolation site (geographic location), and plant host species. The width of each flow is proportional to the number of isolates. Total counts for each category are displayed adjacent to the respective data labels.

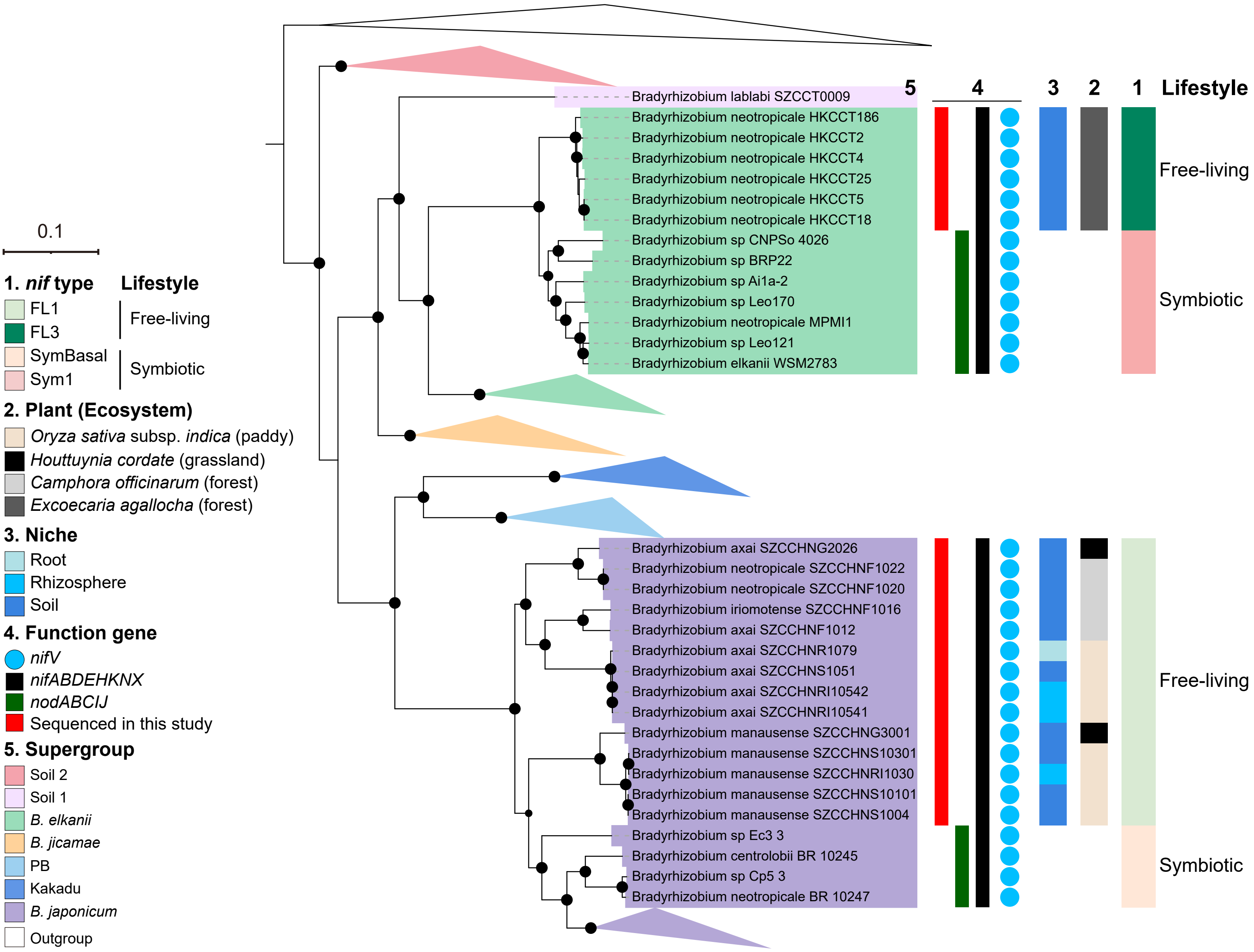


**Figure S3.** Concise phylogeny of the *Bradyrhizobium* emphasizing key representatives from the Soil 1, *B. elkanii*, and *B. japonicum* supergroups, along with associated ecological and genomic traits. Information on other supergroups is not presented in detail. The tree is a simplified version of **Fig. 1A**, with strains from Xanthobacteraceae as the outgroup. Black circles on nodes indicate ultrafast bootstrap values ≥ 95% calculated by IQ-Tree. The scale bar represents 0.1 nucleotide substitution per site.

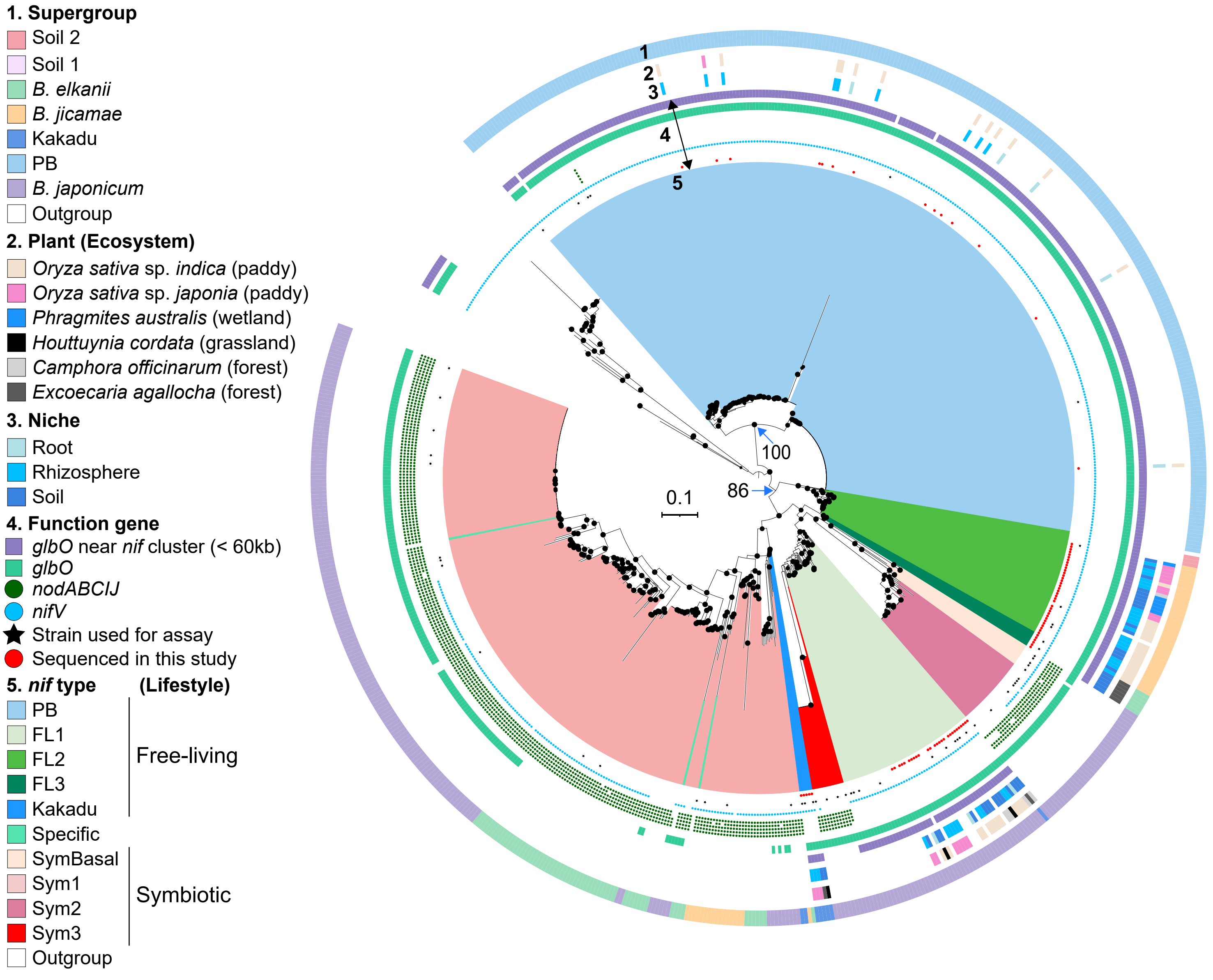


**Figure S4.** *Nif* gene phylogeny of *Bradyrhizobium* based on concatenated alignments of *nifABDEH-KNX*, showing the evolutionary relationships of different nif gene clusters. The tree was constructed under the LG+PMSF(C60)+G+F model, with *nif* genes from *Rhodopseudomonas*, *Azorhizobium*, and *Xanthobacter* as the outgroup. Black circles on nodes indicate ultrafast bootstrap values ≥ 95%, as calculated by IQ-Tree. The bootstrap values for key nodes are also indicated. The scale bar represents 0.1 nucleotide substitutions per site.

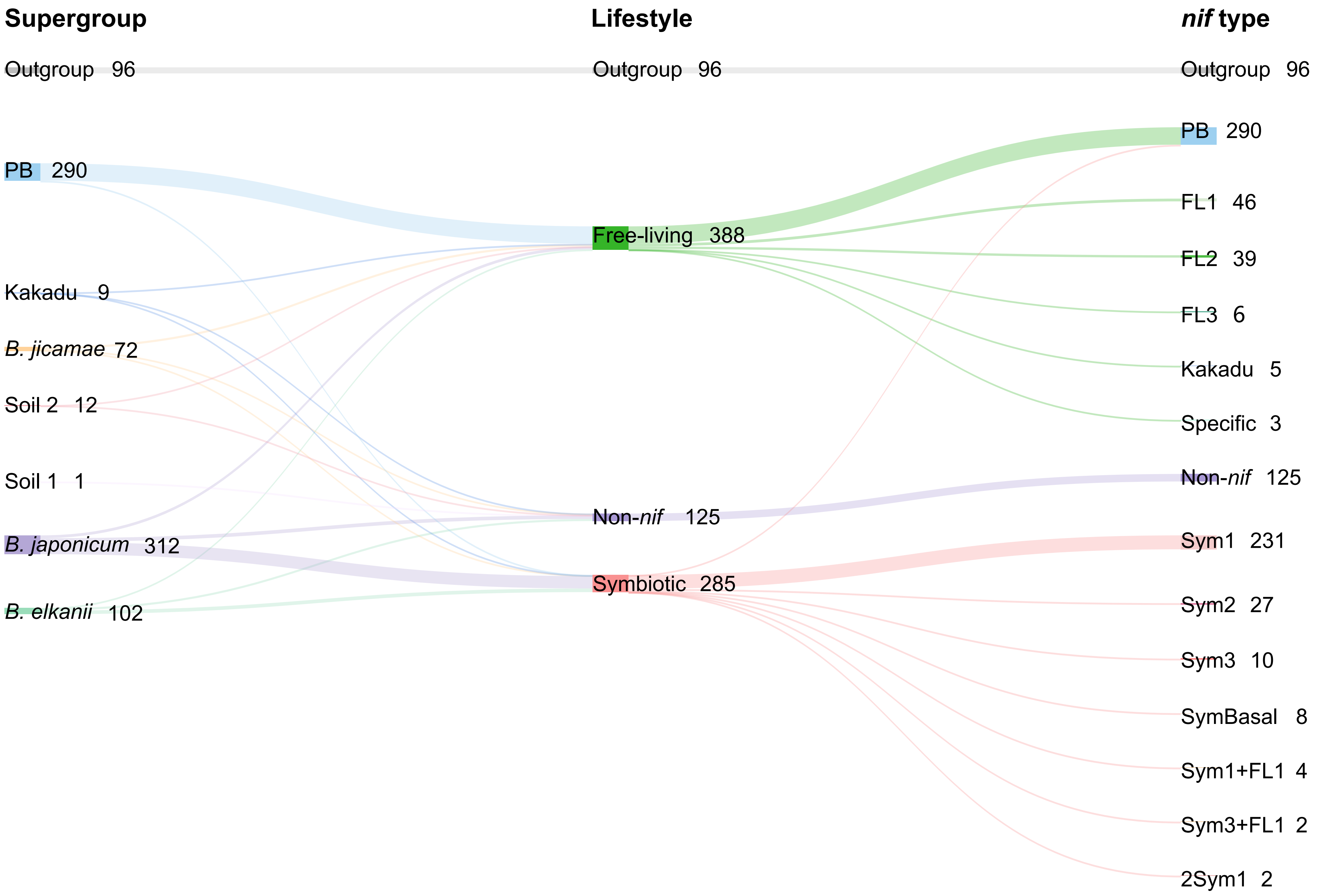


**Figure S5.** Distribution of *Bradyrhizobium* strains (including both newly isolated strains and downloaded genomes) across phylogenetic supergroups, lifestyles, and *nif* types. Sankey diagram illustrates the flow of strain counts from left to right through three categorical levels: supergroup (as defined in **Fig. 1A**), lifestyle (free-living: carrying only *nif* genes, symbiotic: carrying both *nod* and *nif* genes, or Non-*nif*: lacking both *nif* and *nod* genes), and *nif* type (as defined in **Fig. S4**). The width of each flow is proportional to the number of strains. Total counts for each category are displayed adjacent to the respective data labels.

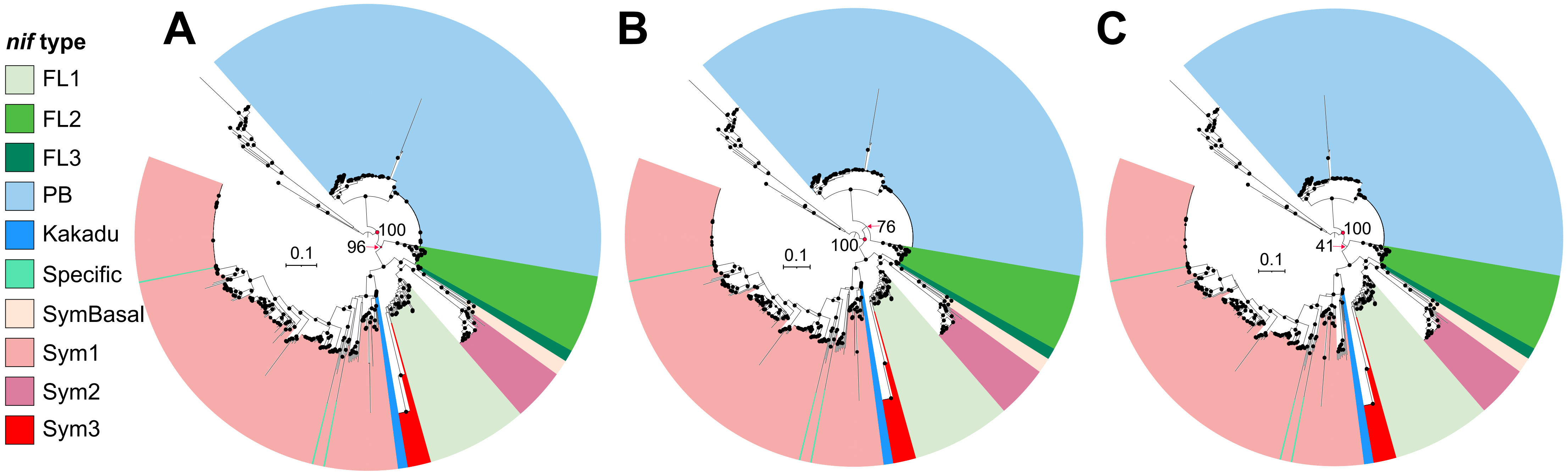


**Figure S6.** *Nif* gene phylogenies of *Bradyrhizobium* inferred under different evolutionary models. The *nif* trees were constructed using the (A) LG+G+F, (B) LG+C20+G+F, and (C) LG+C40+G+F models under the Posterior Mean Site Frequency (PMSF) approximation. The *nif* tree of *Bradyrhizobium* inferred under the LG+C60+G+F model is displayed in **Fig. S4**. Black circles at nodes indicate ultrafast bootstrap values ≥ 95% calculated by IQ-Tree. The horizontal scale bar represents 0.1 nucleotide substitution per site.

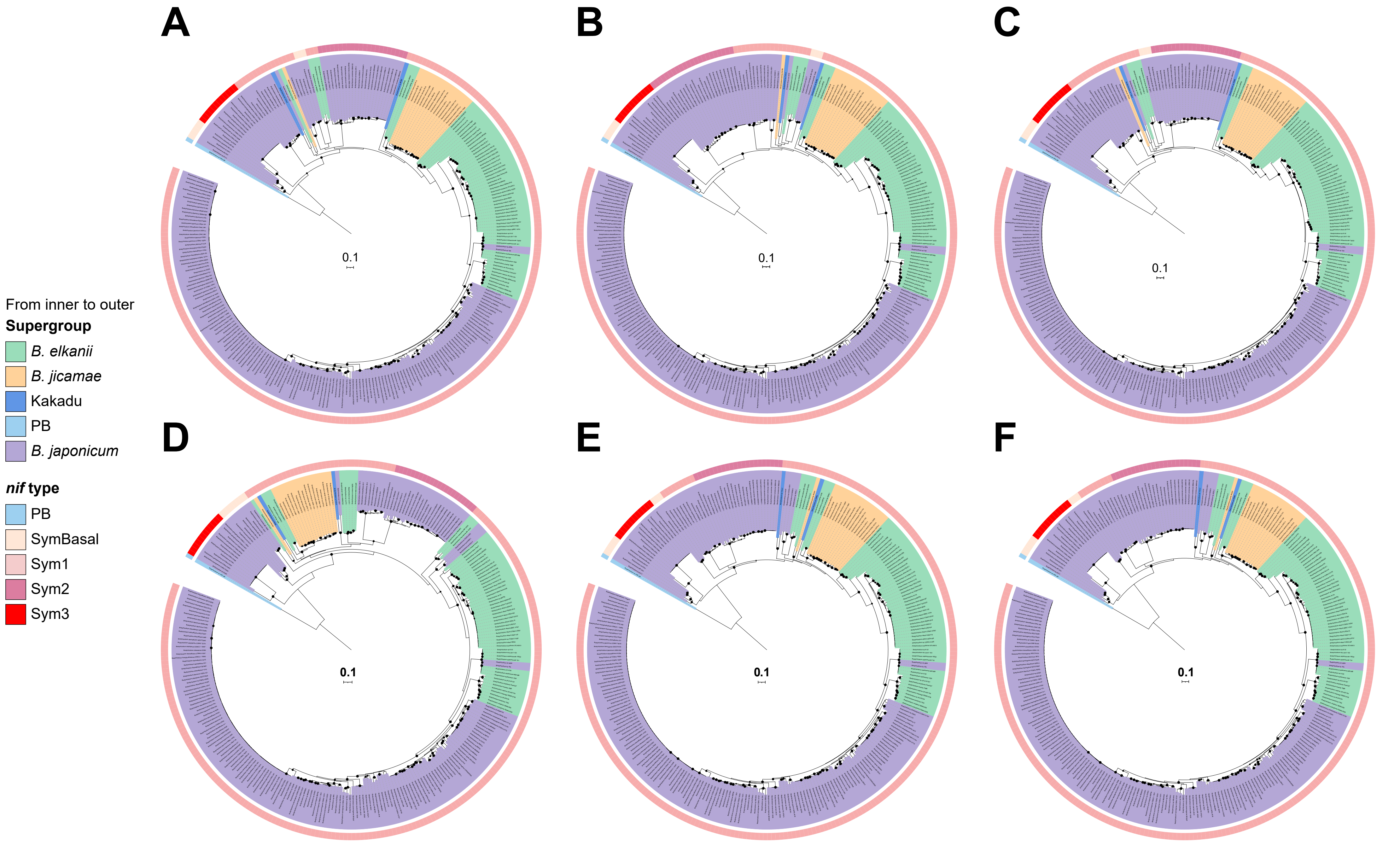


**Figure S7.** *Nod* gene phylogeny of *Bradyrhizobium* inferred from concatenated (A-C) *nodABC* (requiring at least two of the three genes: *nodA*, *nodB*, *nodC*) and (D-F) *nodABCIJ* (requiring at least four of the five genes: *nodA*, *nodB*, *nodC*, *nodI*, *nodJ*) alignments under the following substitution models: (A, D) LG+G+F, (B, E) LG+C20+G+F, and (C, F) LG+C60+G+F under the Posterior Mean Site Frequency (PMSF) approximation. All trees were rooted with the minimum variance (MV) method. Black circles at nodes indicate ultrafast bootstrap values ≥ 95% as calculated by IQ-Tree. The scale bar represents 0.1 substitutions per site.

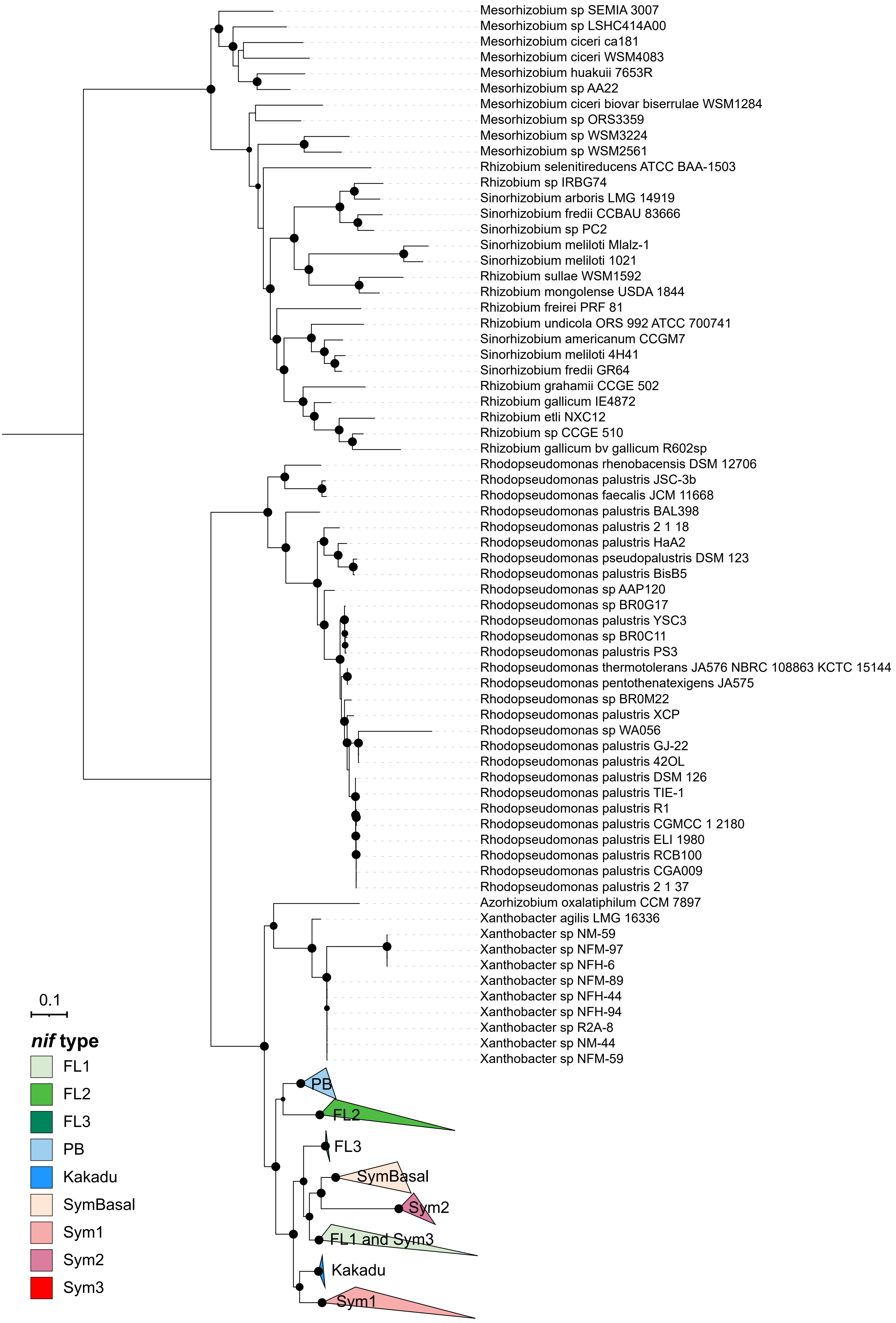


**Figure S8.** *Nif* gene phylogeny of *Bradyrhizobium* including an expanded outgroup. The maximum-likelihood tree was constructed from concatenated alignments of *nifABDEHKNX* under the LG+G+F model. *Nif* genes from *Rhodopseudomonas*, *Azorhizobium*, *Xanthobacter*, *Sinorhizobium*, *Mesorhizobium*, and *Rhizobium* were used as the outgroup. Black circles at nodes indicate ultrafast bootstrap values ≥ 95%, as calculated by IQ-Tree. The following *nif* clusters are marked on the tree: FL1, FL2, FL3, Kakadu, PB, SymBasal, Sym1, Sym2, and Sym3. The scale bar represents 0.1 nucleotide substitutions per site.

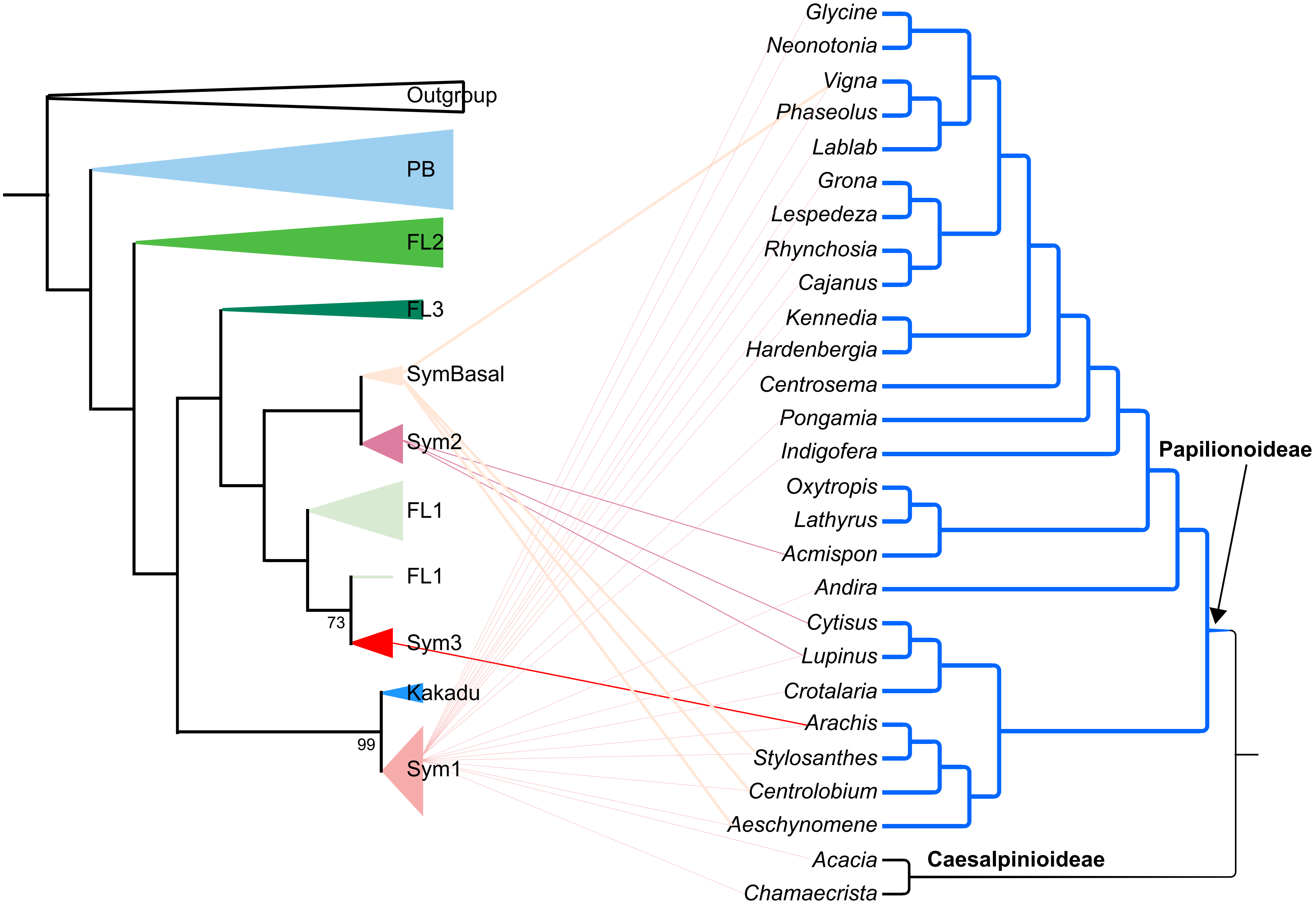


**Figure S9.** Concise view of the phylogenies of different *nif* types in *Bradyrhizobium* (left) and the Fabaceae (right, including representative genera from the subfamilies Papilionoideae and Caesalpinioideae). The left tree is based on **Fig. 1B**, with branch lengths omitted for clarity. The right tree is obtained from https://timetree.org/references (Zhao et al., 2021). Connecting lines between the two trees indicate the original host plant associations of symbiotic *Bradyrhizobium* strains.

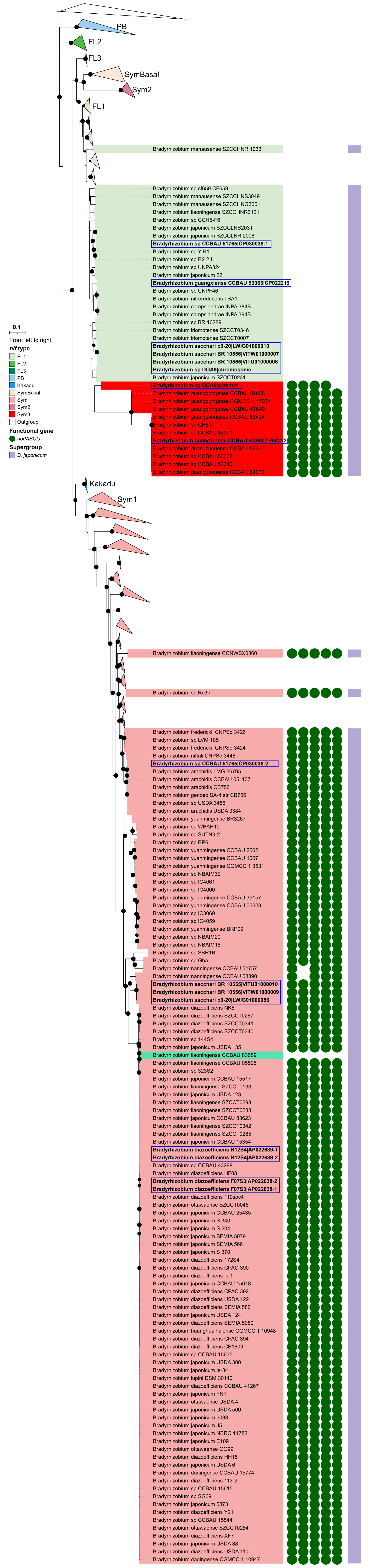


**Figure S10.** Distribution of *Bradyrhizobium* carrying two *nif* clusters in the *nif* phylogeny. This simplified *nif* phylogeny was inferred from **Fig. S4**. The scale bar represents 0.1 nucleotide substitutions per site.

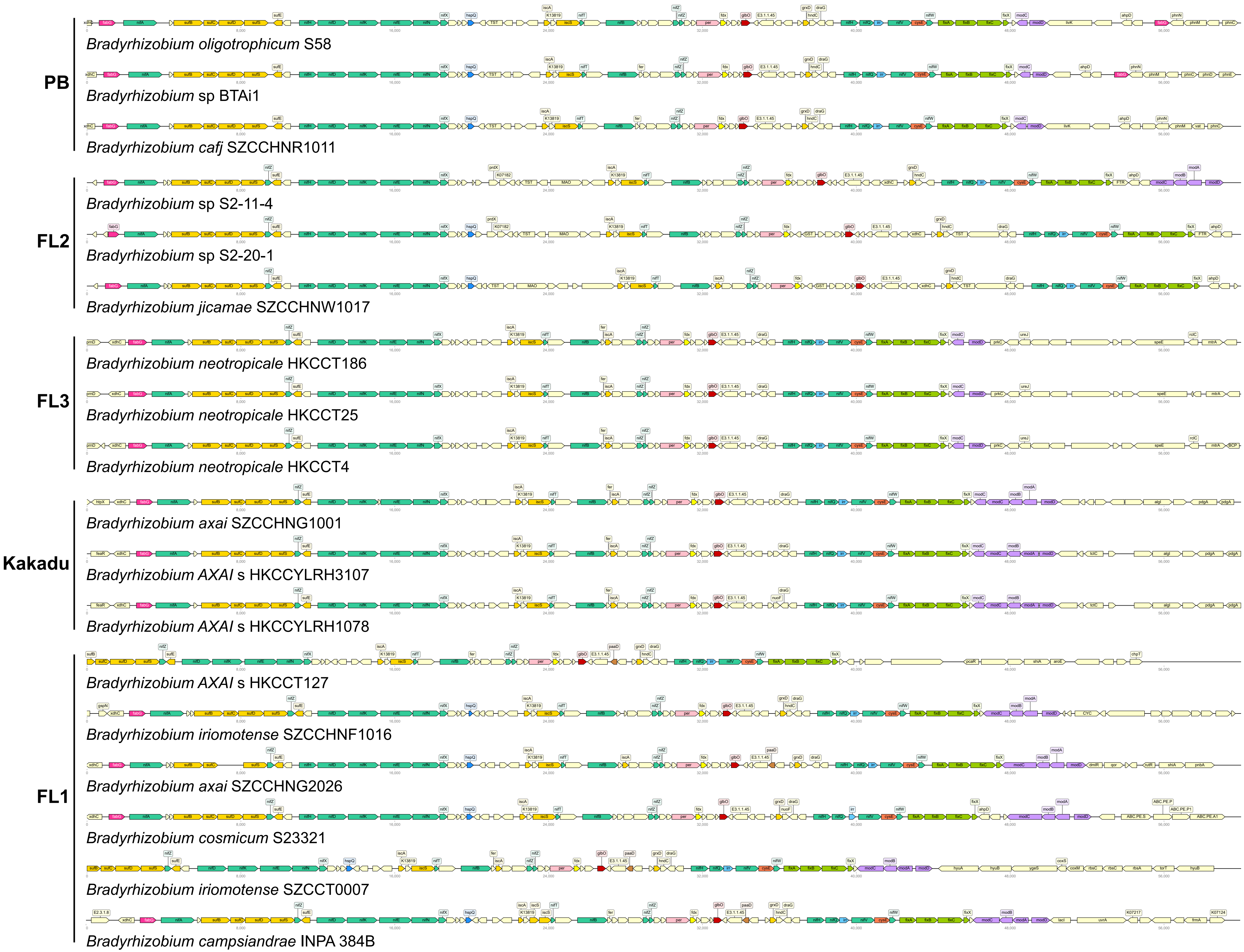


**Figure S11.** Genomic organization of *nif* clusters in representative *Bradyrhizobium* strains capable of free-living nitrogen fixation. Genes with different functional categories are color-coded as indicated.

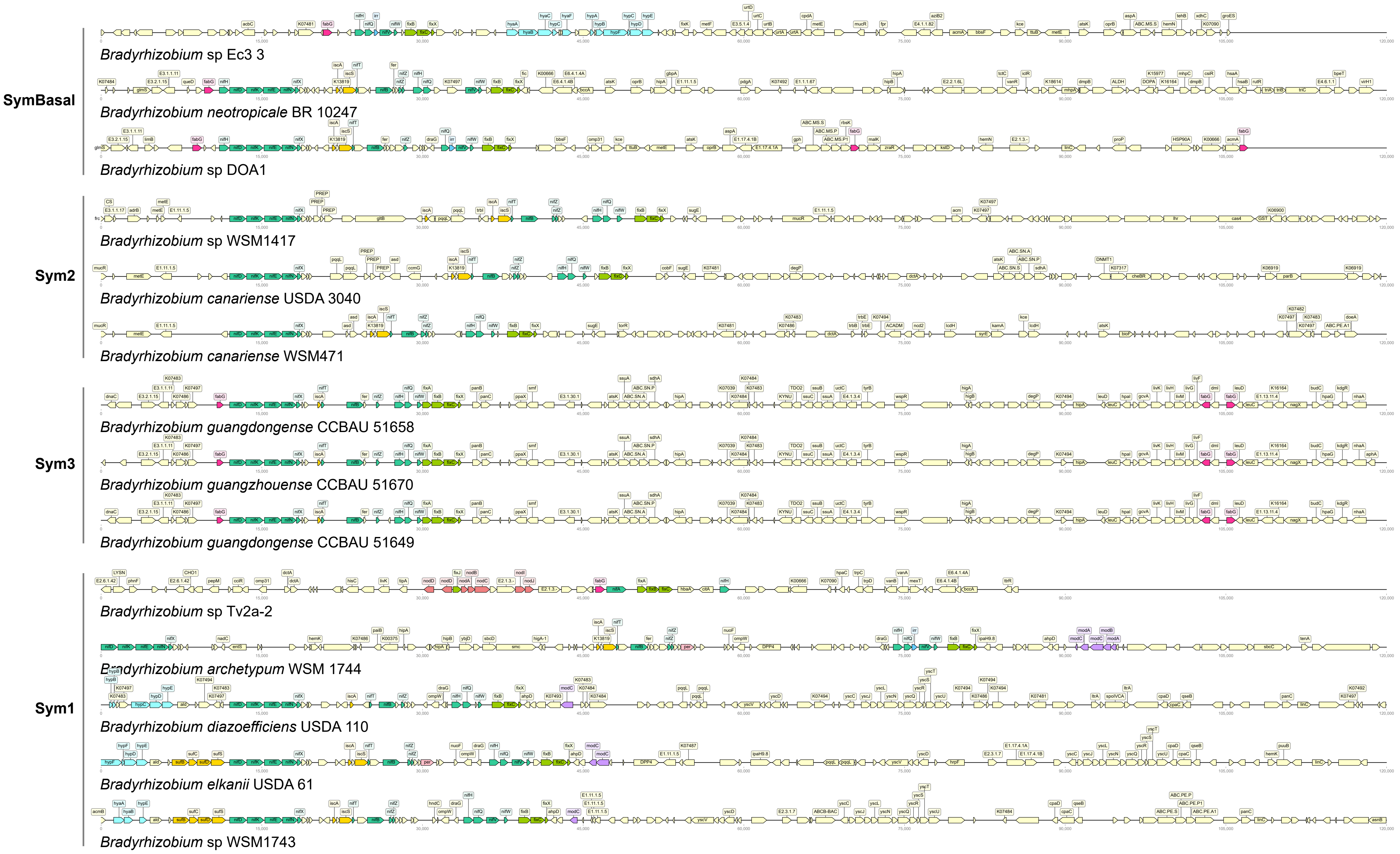


**Figure S12.** Genomic organization of *nif* clusters in representative symbiotic *Bradyrhizobium* strains. Genes with different functional categories are color-coded as indicated.

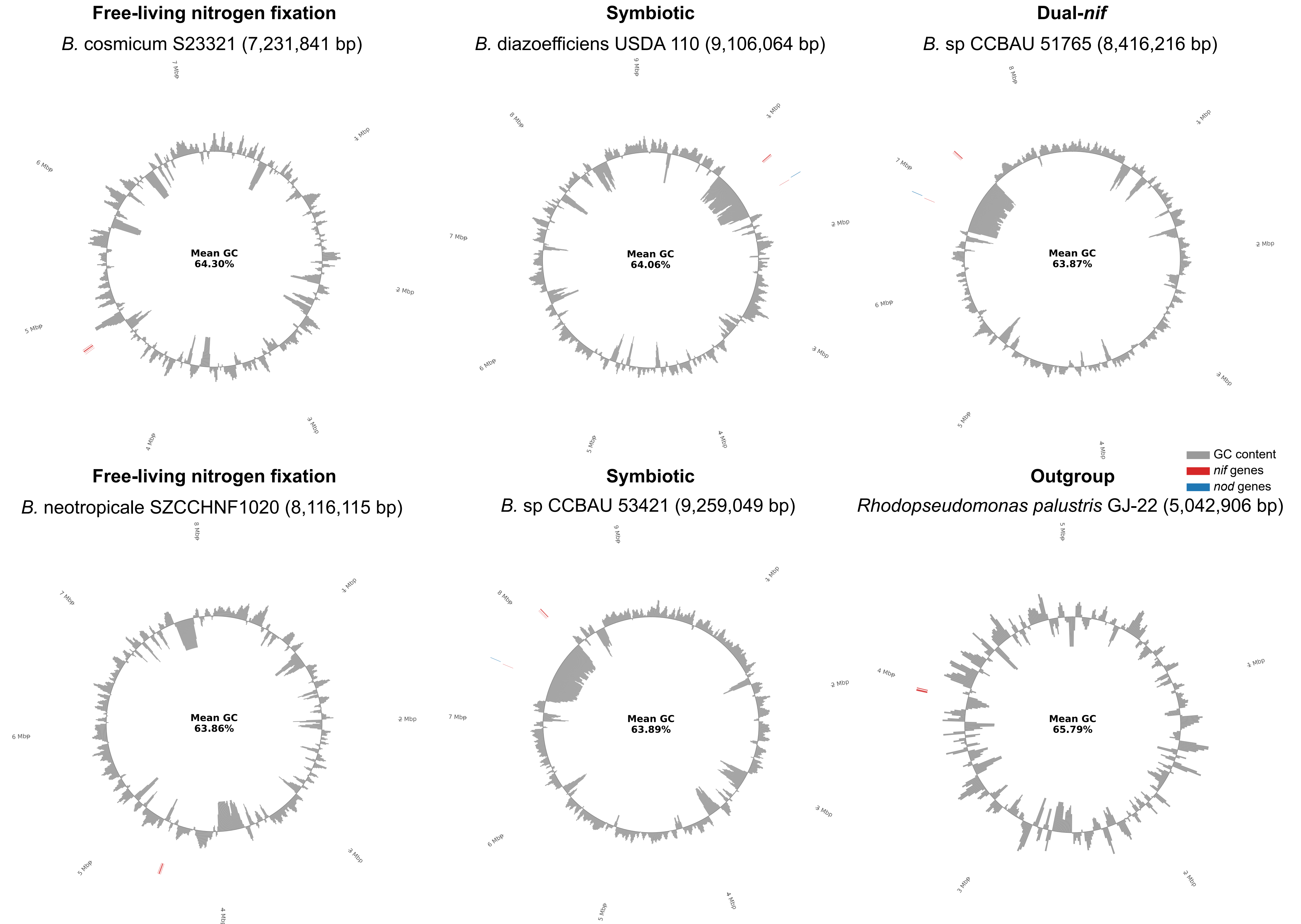


**Figure S13.** Genomic GC content shift in representative *Bradyrhizobium* strains and outgroup members. The plot shows the local GC content deviation (shift) from the genome-wide average baseline, calculated at each genomic position (or within a sliding window). Positive values indicate regions with higher GC content than the genome average, and negative values indicate lower GC content. Data for all free-living and symbiotic complete-genome *Bradyrhizobium* strains are available at: https://figshare.com/s/a657b50b97694bd7373a and https://figshare.com/s/0f6284b4f9df6e53babb, respectively. Data for representative outgroup complete-genome members are available at: https://figshare.com/s/543feda7905ee2d3edc5.

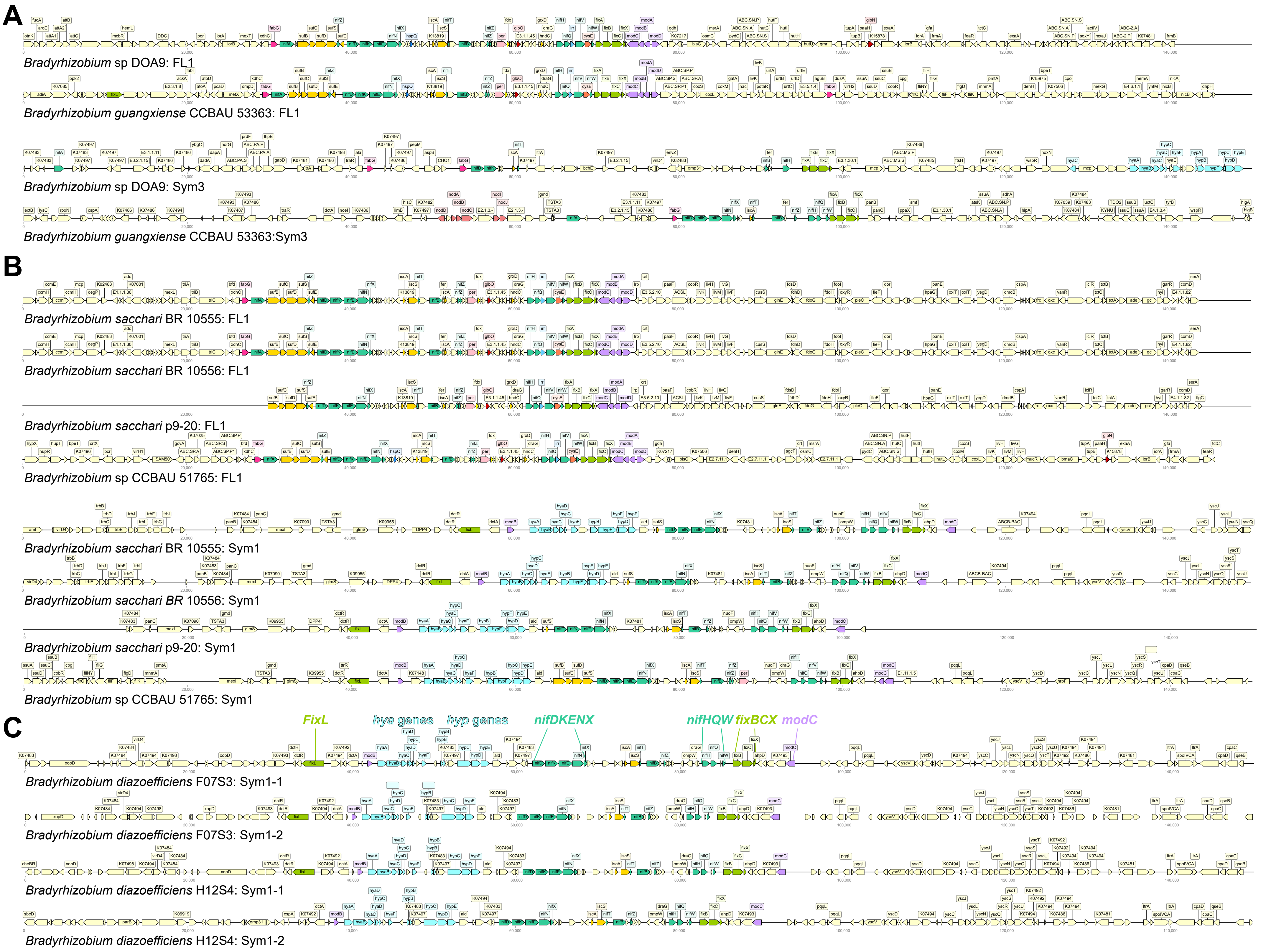


**Figure S14.** Genomic organization of two *nif* clusters in eight *Bradyrhizobium* strains. The arrangement of genes within two *nif* cluster regions present in the same genome. (A) Strains carrying one FL1 and one Sym3 *nif* cluster (Sym3+FL1). (B) Strains carrying one FL1 and one Sym1 *nif* cluster (Sym1+-FL1). (C) Strains carrying two symbiosis islands (designated 2Sym1). Genes with different functional categories are color-coded as indicated.

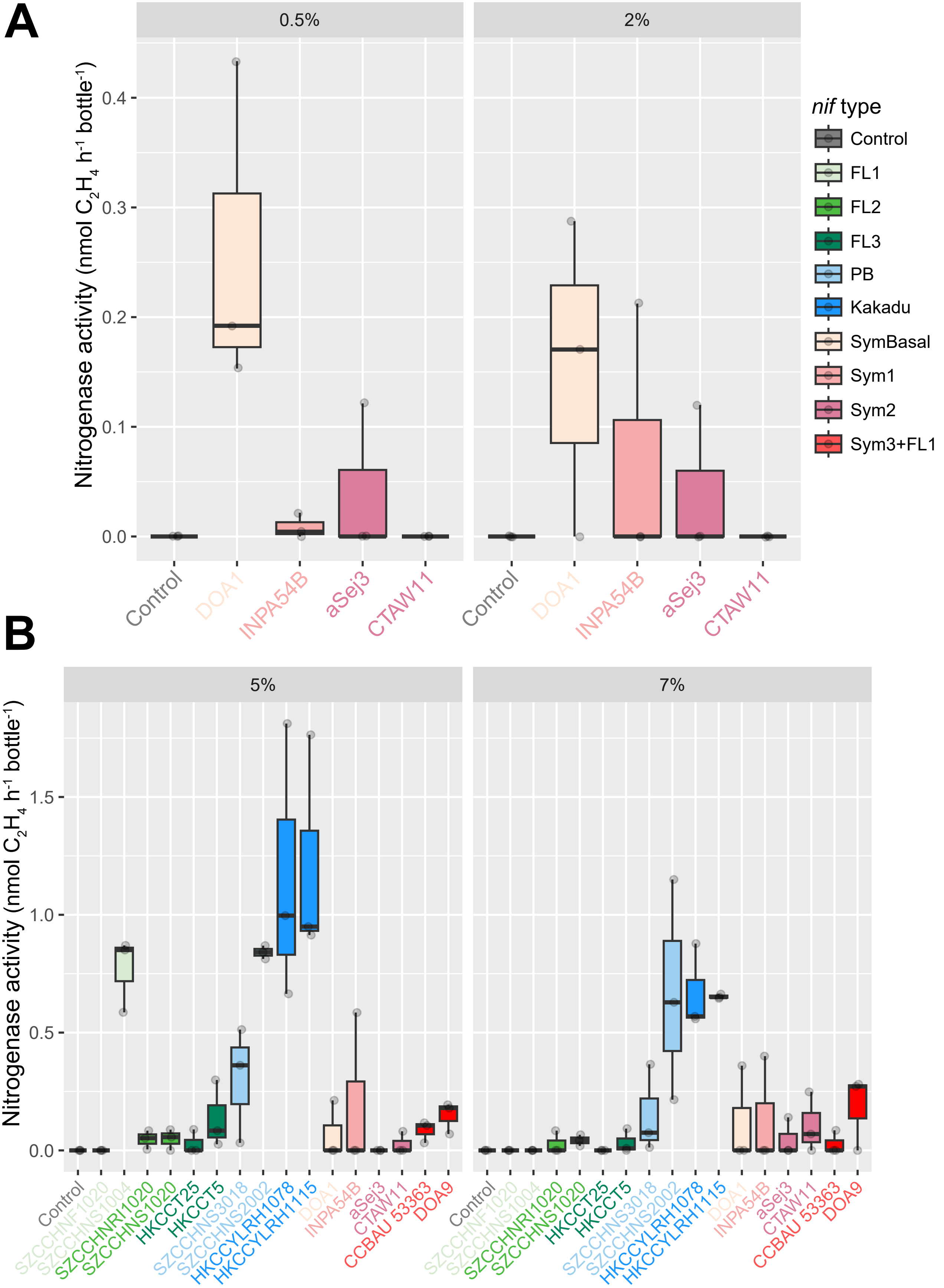


**Figure S15.** Acetylene reduction assay (ARA) measuring ethylene production by *Bradyrhizobium* strains under free-living conditions across different oxygen concentrations. (A) Ethylene production by strains with various symbiotic *nif* types grown in 65 mL vials containing N-free arabinose-gluconate medium with 10% acetylene under 0.5% and 2.0% $O_2$. (B) Ethylene production by strains representing FL1, FL2, FL3, PB, Kakadu, SymBasal, Sym1, Sym2, and Sym3+FL1) under 5.0% and 7.0% $O_2$. All Sym3 strains lacking *nifV* (except dual-*nif* strains) were not selected for the acetylene reduction assay under free-living conditions. Error bars represent standard errors of the mean.

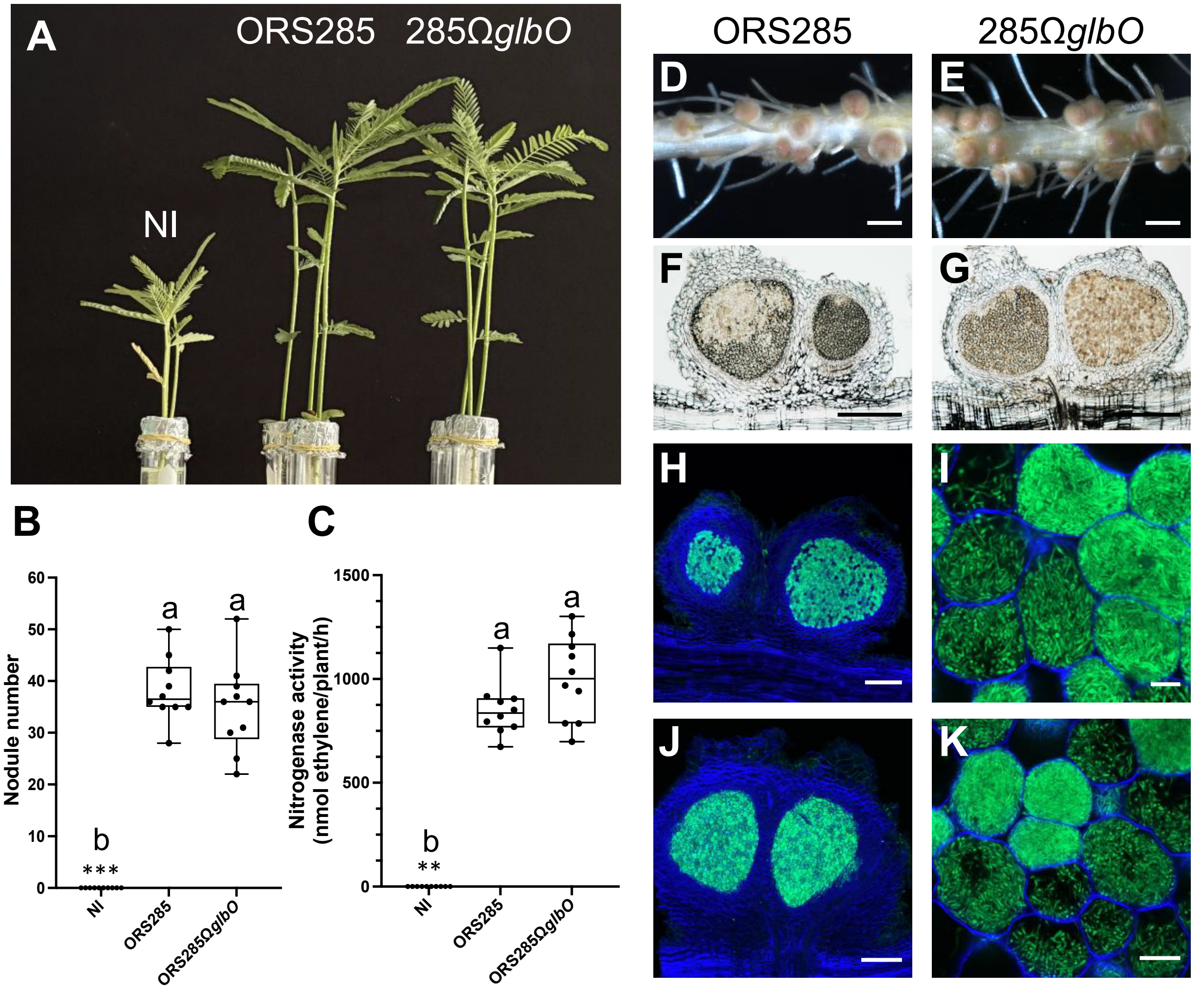


**Figure S16.** The *glbO* mutation has no impact on the symbiotic properties of the ORS285 strain on *A. afraspera*. (A) Photo of *A. afraspera* aerial parts at 17 days post-inoculation (dpi) with *Bradyrhizobium* sp. ORS285 strain and its *glbO* derivative mutant. NI: non-inoculated. (B) and (C) Nodule number and acetylene-reducing activity of *A. afraspera* plants at 17 dpi inoculated with ORS285 strain and its *glbO* derivative mutant. NI: non-inoculated. The box plot show the results obtained on 10 plants. ** $P < 0.005$ , *** $P < 0.0005$, significant differences were observed between the wild type strain and each tested condition using a non-parametric Krustal-Wallis test, NS: not significant. (D and E) Photos of the roots and nodules induced by ORS285 strain and its *glbO* derivative mutant. Scale bars: 0.2 cm. (F and G) Cross section of nodules induced by ORS285 strain and its *glbO* derivative mutant observed using light microscopy showing the internal infected tissue and the peripheral vascularization. Scale bars: 500 µm. (H to K) Confocal microscopy images of a section of nodules induced by ORS285 strain and its *glbO* derivative mutant. The strains were tagged with GFP and calcofluor staining was used to observe the plant cell wall (blue, plant cell wall). Scale bars: H and I, 200 µm; J and K, 10 µm.

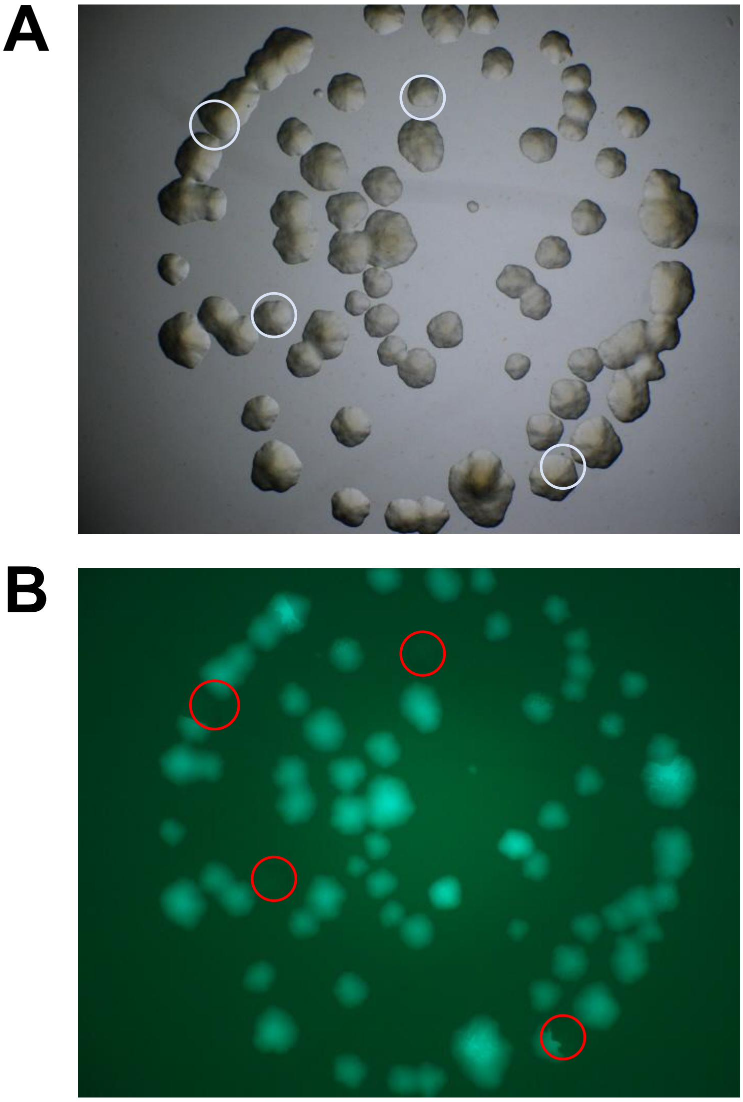


**Figure S17.** Plasmid stability assay in *Bradyrhizobium elkanii* USDA61. Stability of plasmid pVO155-*npt2*-*bjGFP*-*repABC*-C-NK6 was assessed in strain USDA61. After one week of cultivation of a plasmid-harboring clone in YM medium without antibiotic selection, serial dilutions were spotted onto YM agar plates without antibiotics. After 6 days of incubation, colonies within each spot were counted under white and green light. Colonies lacking GFP fluorescence, indicating plasmid loss, are outlined in red.
