## Supplementary material for "The free-living wellspring of symbiotic nitrogen fixation in *Bradyrhizobium*": SI files

**This PDF file includes:**

**Supplementary Text:** Methods

**Supplementary references**

**Figures S1 to S17**

### Supplementary Text: Methods

<u>Soil and plant sampling and processing</u>

To substantially expand the genomic representation of free-living nitrogen-fixing (FL*nif*) *Bradyrhizobium*, we conducted extensive sampling across major ecosystems in China, targeting a diverse array of non-legume plants and adjacent soils (0 - 5 cm depth). Sampling encompassed five geographically distinct regions (**Fig. S1**) with significant variations in precipitation and temperature to capture broad ecological diversity. Cropland samples included rice (*Oryza sativa* from Hunan, Liaoning, and Hong Kong) and maize (*Zea mays* from Shanxi and Anhui provinces). From natural grassland, forest, and wetland ecosystems, we collected *Houttuynia cordata*, *Camphora officinarum*, and *Phragmites australis* in Hunan, and *Excoecaria agallocha* in Hong Kong. All samples were transported to the laboratory in sterile bags within iceboxes for further processing.

We processed three distinct niches for each sample, which were classified as follows: soil samples collected far from plant roots were stored as bulk soil. Rhizosphere samples were obtained by removing loose soil attached to roots and then vigorously vortexing root segments in phosphate-buffered saline to dislodge tightly attached soil particles, followed by centrifugation, as previously described (1). The roots were then thoroughly washed to remove all residual soil and designated as endosphere samples.

<u>Isolation and identification of *Bradyrhizobium* strains</u>

Each root, rhizosphere, and bulk soil sample were separated into two portions for subsequent processing. One portion was subjected to a four-day enrichment under controlled light and temperature conditions (28°C with 70% relative humidity on a 16:8-h day: night cycle) to selectively promote the growth of Photosynthetic *Bradyrhizobium* supergroup strains. The other portion (non-enriched) was used as the original sample to collect other *Bradyrhizobium* supergroup strains. Both sample portions (enriched and non-enriched) were used for *Bradyrhizobium* isolation on selective media according to our previous study (2). Taxonomic classification of each isolate was determined based on their 16S rRNA gene sequence. *Bradyrhizobium* isolation from maize samples (Shanxi and Anhui provinces) had limited success.

Analysis of basic soil characteristics

Soil samples were processed to measure the basic soil characteristics upon arrival at the laboratory. Soil water content was determined by oven-drying 10.00 g of fresh soil at 105°C for 6 hours. The weight loss was divided by the dry weight to calculate the soil water content after weighing the dried soil samples. Soil pH was measured using a pH electrode with a soil-to-deionized water ratio of 1:5 (w/w). Soil organic carbon (SOC) was analyzed using the potassium dichromate oxidation process combined with the heating method. Total nitrogen (TN) was determined via the Kjeldah method (3), and total phosphorus (TP) was measured using the sodium hydroxide melting-molybdenum antimony colorimetric method (3). Available phosphorus (AP) was ascertained using the molybdenum antimony anti-colorimetric method (3). Soil inorganic nitrogen fractions, including nitrite nitrogen ($NO_2^-$), nitrate nitrogen ($NO_3^-$), and ammonium nitrogen ($NH_4^+$), were extracted using by 2.0 mol $L^{-1}$ potassium chloride (KCl) solution and detected by continuous flow analysis (4). Dissolved organic carbon and nitrogen (DOC and DON) were extracted by adding 25 mL of Milli-Q water to 5 g of fresh soil, followed by shaking at 220 rpm for 30 minutes. The supernatant was filtered through a 0.45 μm quantitative filter membrane and then analyzed using a TOC/TN analyzer (Shimadzu, Analytical Sciences, Kyoto, Japan) (5, 6). Microbial biomass carbon and nitrogen (MBC and MBN) were determined using the chloroform fumigation-extraction method (5, 6). MBC and MBN were calculated as the difference in DOC and DON concentration between the fumigated and unfumigated soil samples and multiplied by a factor of 2.22. Heavy metal concentrations, including iron (Fe) and molybdenum (Mo), were measured using inductively coupled plasma mass spectrometry (ICP-MS). All sample-associated metadata are provided in Supplementary **Dataset S1**.

Genome sequencing, assembly, and annotation

Genomic DNA from each of the 88 novel isolates was extracted using the TIANamp Genomic DNA Kit (TIANGEN) according to the manufacturer's protocol. The DNA quality and quantity meeting the required standards (A260/A280 > 1.8, A260/A230 > 2.0, and A260/A270 > 1.0) were sent to Wuhan Huada Gene Biotechnology Company for whole genome sequencing.

The untrimmed adapters conjunct with raw reads were identified by BBMerge implemented in BBmap package v38.79 (7). The adapters and low-quality reads were trimmed using

Trimmomatic v0.39 (8) and reads shorter than 40 bp were discarded. The quality of the remaining reads was assessed using FastQC (https://www.bioinformatics.babraham.ac.uk/projects/fastqc/) (9). The remaining paired-end reads with high quality were assembled using SPAdes v3.10.1 with default parameters (10). Those contigs with lengths greater than 1,000 bp and k-mer coverage over five were retained for further analysis. The 88 high-quality genome assemblies together with the downloaded 807 genomes were predicted using Prokka v1.12 (11) and annotated using the KEGG database (12).

Phylogenomic and *nif* gene tree construction

To identify the phylogenetic distribution of the isolates in the *Bradyrhizobium*, a phylogenomic tree of *Bradyrhizobium* (**Fig. 1A**) was built using IQ-Tree v2.2.0 (13) with 88 isolates and 807 public genomes (including outgroup members) downloaded from the NCBI Genbank database based on 123 shared single-copy genes (14). The following parameter "-s alignment -spp partition -m MFP -mset LG,WAG,JTT -mrate E,G,I,G+I -bb 1000" was applied so that each gene was allowed to automatically select its best-fit substitution model by ModelFinder implemented in IQ-Tree (13). Branch support was evaluated using 1000 ultrafast bootstrap approximations (15).

To specifically investigate the evolutionary history of nitrogen fixation, the *nif* gene tree (**Fig. S4, S6**) was constructed based on the concatenated amino acid sequences of core *nif* genes (*nifABDEHKNX*) with the parameter "-s alignment -spp partition -m MFP -mset LG -mrate E,G,I,G+I -bb 1000". These *nif* genes were selected for their central role in nitrogenase function and high evolutionary conservation (14, 16). *Bradyrhizobium* strains were systematically classified into three ecological types based on the presence or absence of the *nod* and *nif* gene clusters (**Fig. S5, Dataset S1, S2**). Strains carrying both *nifABDEHKNX* and *nodABCIJ* genes (allowing a maximum of two missing genes) were categorized as *nod*-carrying (symbiotic) members. Strains carrying only *nifABDEHKNX* genes were classified as free-living *nif*-carrying (FL*nif*) members, among which *nod*-free but nodulating members (traditionally called Photosynthetic *Bradyrhizobium* supergroup or PB supergroup) were a special type that lack *nod* genes but can form nodules with *Aeschynomene* (2, 17). Strains lacking both *nod* and *nif* genes were grouped as free-living non-*nif* carrying (FLnon*nif*) members (**Fig. S5, Dataset S1, S2**).

Construction of a *glbO* insertion mutant in *Bradyrhizobium* sp. ORS285

To investigate the functional role of *glbO*, which encodes a putative truncated hemoglobin protein located in the vicinity of the *nif* genes in FL*nif Bradyrhizobium*, an insertional mutant of *glbO* (BRAO285v2_1830) was constructed in ORS285 strain. The downstream gene of *glbO* is in the opposite direction, eliminating the risk of polar mutation. The insertional mutant (ORS285Ω*glbO*) was obtained by single-crossing over. A 233-bp internal fragment of the target gene was amplified by polymerase chain reaction (PCR) using the primers (285.*glbO*.in.f/ 285.*glbO*.in.r) (**Dataset S9**) and cloned into a derivative of the non-replicative pVO155-*Sm*-P*npt2-GFP*, which carries a kanamycin and a streptomycin resistance genes and a constitutive GFP under the *npt2* promoter (P*npt2*) (18, 19). The resulting plasmid was then transferred into ORS285 by conjugation (20) and mutants were selected on YM (21, 22) antibiotic-containing plates (Cefotaxime 20 μg/mL, Kanamycin 50 μg/mL and Spectinomycin 20 μg/mL). All cloning steps were performed using standard molecular biology techniques.

Introduction of *glbO* into *Bradyrhizobium elkanii* USDA61

To construct the shuttle plasmid pVO155-P*npt2-bjGFP-repABC*-C-NK6, the plasmid pVO155-P*npt2-bjGFP* was first generated by amplifying the P*npt2-bjGFP* region from plasmid pRJPaph-bjGFP (23) using primers pVO155-P*npt2-bjGFP*.f and pVO155-P*npt2-bjGFP*.r (**Dataset S9**). The PCR product was digested with KpnI and cloned into the corresponding site of pVO155 (24). The *repABC* region from plasmid pNK6c of *Bradyrhizobium diazoefficiens* NK6 was then amplified using primers RepABC-NK6-pC.f and RepABC-NK6-pC.r (**Dataset S9**) and cloned into the BamH1/XbaI sites of the pVO155-P*npt2-bjGFP* vector. The resulting plasmid, pVO155-P*npt2-bjGFP-repABC*-C-NK6, was introduced into *Bradyrhizobium elkanii* USDA61 by electroporation as previously described (25).

Plasmid stability was assessed using three independent transformants grown in YM medium without antibiotic selection for one week, followed by plating onto YM agar without antibiotics. After five days of incubation, fluorescence microscopy revealed that more than 95% of the colonies exhibited GFP fluorescence (**Fig. S17**). We additionally assessed plasmid retention based on maintenance of the plasmid-associated kanamycin resistance marker. Among the 500 colonies tested, all GFP-positive colonies retained kanamycin resistance (i.e., were able to grow in the presence of kanamycin at 50 μg/mL), whereas all GFP-negative colonies were sensitive to kanamycin. Collectively, these results demonstrate that pVO155-P*npt2-bjGFP-repABC*-C-NK6 is highly stable in the USDA61 strain under non-selective growth conditions.

Following confirmation of plasmid stability, the *glbO* gene was introduced under the control of the constitutive *npt2* promoter. The *npt2* promoter and the *glbO* gene from strain ORS278 were fused by overlapping PCR and cloned into the SpeI restriction site of pVO155-P*npt2-bjGFP-repABC*-C-NK6. The resulting plasmid, pVO155-P*npt2-bjGFP-repABC*-C-P*npt2-glbO*, as well as the corresponding empty vector control, were independently introduced into *B. elkanii* USDA61 by electroporation as described previously (25). All constructs generated in this study, including primers and cloning strategies, are listed in **Dataset S9**.

<u>Acetylene reduction assay (ARA)</u>

To systematically compare the nitrogen-fixing ability in different *nif* type *Bradyrhizobium*, several representative strains from each *nif* cluster were selected (**Fig. 3A, Dataset S1, S2**) for an acetylene reduction assay under a semi-solid buffered nodulation medium (BNM) without oxygen control. For this experiment, the bacteria were grown in 9 mL vacuette® tubes (Greiner Bio-One GmbH) containing 2 mL of BNM with 0.8% agar (semi-solid) and supplemented with 10 mM succinate and 10 mM arabinose as the carbon source. The tubes were incubated in a dark environment at 28 °C for 6 days (26). At the beginning of the experiment, the headspace was replaced with 10% (v/v) acetylene. After six-day incubation, nitrogenase activity was determined by measuring the amount of ethylene produced by the bacterial culture using gas chromatography (21).

To systematically compare the oxygen tolerance of nitrogenase of different *nif* type *Bradyrhizobium*, two representative strains from each *nif* cluster were selected (**Fig. 3B**, **Dataset S1, S2**) for an acetylene reduction assay under different $O_2$ concentrations (0.5%, 2%, 5%, and 7%). As previous studies have shown that *Bradyrhizobium* can fix nitrogen under free-living conditions at $O_2$ concentrations below 5% (27), a gradient of 0.5%, 2%, and 5% $O_2$ was chosen for the ARA to systematically compare the nitrogen fixation efficiency among different *nif* type *Bradyrhizobium*. An additional 7% $O_2$ concentration was included to investigate whether nitrogen fixation could be sustained at supra-threshold oxygen levels (> 5%).

The nitrogenase activity of various *nif* type *Bradyrhizobium* strains was characterized in 65 mL tubes containing 20 mL of nitrogen-free modified arabinose-gluconate (MAG) liquid medium. The strains were initially grown in 14 mL test tubes containing 5 mL of nitrogen-free MAG liquid medium (14). The tubes were then incubated at 28°C on a rotary shaker at 170 rpm for 4 days to generate seed cultures. The bacterial cells were then harvested by centrifugation at 12,000 rpm for 2 minutes at 4°C, and the resulting pellets were washed three times with sterile Milli-Q water. The optical density ($OD_{600}$) of the bacterial suspensions was normalized to 0.2 (28). Subsequently, 100 µL of each suspension was aseptically inoculated into 65 mL sterile bottles containing 20 mL of fresh, nitrogen-free MAG liquid medium using an injection syringe, prepared without $NH_4Cl$ to ensure nitrogen-limiting conditions.

The inoculated bottles were incubated at 28°C with a continuous shaking speed of 170 rpm for 4 days under $O_2$ concentrations of 0.5%, 2.0%, 5.0% and 7%, with $N_2$ making up the balance. To maintain the precise oxygen concentrations, the headspace gas in each bottle was replaced daily with the corresponding gas mixture. For the final 24 hours of incubation, acetylene (10% v/v) was injected into each bottle to initiate the assay (29, 30). After 24 hours, a 1 mL gas sample was extracted from the headspace of each bottle. The ethylene content of these samples was quantified using a gas chromatography (Gas Chromatograph 163, Hitachi, Tokyo, Japan) equipped with an HP Plot Q column. The ethylene production rate per unit time and per unit bottle was calculated in nmol $C_2H_4$ bottle$^{-1}$ h$^{-1}$. The nitrogenase activity of *Bradyrhizobium* was expressed as the ethylene production rate in nmol $C_2H_4$ bottle$^{-1}$ h$^{-1}$.

To compare the nitrogenase activity of ORS285 and its mutant (ORS285Ω*glbO*) under different free-living conditions, we performed ARA in both semi-solid and liquid BNM media, with experimental conditions consistent with **Fig. 3A**. We also performed the ARA for all the complementation strains in semi-solid buffered nodulation medium (BNM).

Kinetics study of ethylene formation

In kinetic studies, the absorbance (at 600 nm) of the bacterial culture was measured at various time intervals by taking 1 mL samples from liquid BNM. The amount of ethylene produced by the bacterial culture at different time points was also measured by injecting a 1 mL gas sample into a gas chromatograph.

Symbiotic analysis on *Aeschynomene indica* and *Aeschynomene afraspera*

The ORS285 strain and its *glbO* mutant derivative used for symbiotic analysis were grown in arabinose-gluconate (AG) medium (31) at 28°C on Petri dishes for one week. Bacteria were harvested from the plate and resuspended in 10 mL of sterile water and the $OD_{600}$ was adjusted to 1.0. *A. indica* and *A. afraspera* plants were cultured as previously described (18). Each strain was inoculated into ten plants with 1 mL of bacterial suspension and the symbiotic properties (number of nodules per plant and nitrogenase enzyme activity) were analyzed 17 days after inoculation (32). Cytological analysis of nodules elicited by ORS285 and ORS285Ω*glbO* were performed as described (33).

<u>Statistical analysis</u>

Nitrogenase activities, measured by the acetylene reduction assay were analyzed in R version 4.5.1 (R core team 2025) using the packages "ggplot2" (34), "lme4" (35), and "emmeans" (https://CRAN.R-project.org/package=emmeans). To account for the phylogenetic non-independence of *Bradyrhizobium* isolates and confounding effects of their evolutionary origins on nitrogenase activity, we quantified the phylogenetic signal of nitrogenase activity across the species tree using Pagel's lambda with the phylosig function (36). If high phylogenetic conservatism was observed, the phylogenetic analysis of variance (PhylANOVA) was performed using the "phylANOVA" function in the phytools (v2.0) package of R (36). The model included a fixed factor (i.e., symbiotic vs. capable of free-living nitrogen fixation) with nitrogenase activity as the primary predictor. The analysis was conducted with 10,000 simulations to generate a null distribution of the F-statistics under the Brownian motion evolution model. The pruned phylogenomic tree of the isolates (53 members for **Fig. 3A** and 16 for **Fig. 3B**) used in the acetylene reduction assay was implemented in the PhylANOVA analysis. To assess the nitrogenase activity variations across *nif* types (**Fig. 3A, B, Dataset S3, S4**), we fitted a linear mixed-effects model (LMM) defining the logarithm-transformed "ARA" values as the responsible variables (35). This model included lifestyles and *nif* types as two fixed effects and strain identity and biological replicate as two random effects. The normal distribution of the logarithm-transformed data was confirmed. The significance of the fixed effects was assessed via two-way ANOVA, with post-hoc pairwise comparisons performed using Tukey's method. To compare the significant differences between the wild-type and mutant strains of ORS285 and USDA61, one-way ANOVA test (**Fig. 4A, B, E**) were performed followed by Tukey's post-hoc analysis.

living growth. *Front Microbiol* 14:1131860.

26. Nouwen N*, et al.* (2017) The role of rhizobial (*NifV*) and plant (*FEN1*) homocitrate synthases in *Aeschynomene*/photosynthetic *Bradyrhizobium* symbiosis. *Sci Rep* 7(1):448.
27. Terakado-Tonooka J, Fujihara S, & Ohwaki Y (2013) Possible contribution of *Bradyrhizobium* on nitrogen fixation in sweet potatoes. *Plant and Soil* 367(1-2):639-650.
28. Siqueira AF, Sugawara M, Arashida H, Minamisawa K, & Sánchez C (2020) Levels of periplasmic nitrate reductase during denitrification are lower in *Bradyrhizobium japonicum* than in *Bradyrhizobium diazoefficiens*. *Microbes and environments* 35(3):ME19129.
29. Mbai F, Magiri E, Matiru V, Nganga J, & Nyambati V (2013) Isolation and characterization of bacterial root endophytes with potential to enhance plant growth from Kenyan Basmati rice. *American International Journal of Contemporary Research* (3): 25.
30. Montes-Luz B*, et al.* (2023) Acetylene reduction assay: A measure of nitrogenase activity in plants and bacteria. *Curr Protoc* 3(5):e766.
31. Sadowsky MJ, Tully RE, Cregan PB, & Keyser HH (1987) Genetic diversity in *Bradyrhizobium japonicum* serogroup 123 and its relation to genotype-specific nodulation of soybean. *Appl Environ Microbiol* 53(11):2624-2630.
32. Bonaldi K*, et al.* (2010) Large-scale transposon mutagenesis of photosynthetic *Bradyrhizobium* sp. strain ORS278 reveals new genetic loci putatively important for nod-independent symbiosis with *Aeschynomene indica*. *Mol Plant Microbe Interact* 23(6):760-770.
33. Songwattana P*, et al.* (2021) Identification of type III effectors modulating the symbiotic properties of *Bradyrhizobium* vignae strain ORS3257 with various *Vigna* species. *Sci Rep* 11(1):4874.
34. Valero-Mora PM (2010) ggplot2: Elegant graphics for data analysis. *Journal of Statistical Software* 35(Book Review 1):1 - 3.
35. Bates D, Mächler M, Bolker BM, & Walker SC (2015) Fitting linear mixed-effects models using lme4. *Journal of Statistical Software* 67(1):1-48.
36. Revell LJ (2024) phytools 2.0: an updated R ecosystem for phylogenetic comparative methods (and other things). *PeerJ* 12:e16505.
